# Transition to invasive breast cancer is associated with progressive changes in the structure and composition of tumor stroma

**DOI:** 10.1101/2021.01.05.425362

**Authors:** Tyler Risom, David R Glass, Candace C Liu, Belén Rivero-Gutiérrez, Alex Baranski, Erin F McCaffrey, Noah F Greenwald, Adam Kagel, Siri H Strand, Sushama Varma, Alex Kong, Leeat Keren, Sucheta Srivastava, Chunfang Zhu, Zumana Khair, Deborah J Veis, Katherine Deschryver, Sujay Vennam, Carlo Maley, E Shelley Hwang, Jefferey R Marks, Sean C Bendall, Graham A Colditz, Robert B West, Michael Angelo

## Abstract

Ductal carcinoma *in situ* (DCIS) is a pre-invasive lesion that is thought to be a precursor to invasive breast cancer (IBC). To understand how the tumor microenvironment (TME) changes with transition to IBC, we used Multiplexed Ion Beam Imaging by time of flight (MIBI-TOF) and a 37-plex antibody staining panel to analyze 140 clinically annotated surgical resections covering the full spectrum of breast cancer progression. We compared normal, DCIS, and IBC tissues using machine learning tools for multiplexed cell segmentation, pixel-based clustering, and object morphometrics. Transition from DCIS to IBC was found to occur along a trajectory marked by coordinated shifts in location and function of myoepithelium, fibroblasts, and infiltrating immune cells in the surrounding stroma. Taken together, this comprehensive study within the HTAN Breast PreCancer Atlas offers insight into the etiologies of DCIS, its transition to IBC, and emphasizes the importance of the TME stroma in promoting these processes.

## Introduction

Ductal Carcinoma in situ (DCIS) is a preinvasive lesion where tumor cells within the breast duct are isolated from the surrounding stroma by a near-continuous layer of myoepithelium and basement membrane proteins. This histologic feature is the central property that distinguishes it from invasive breast cancer (IBC), where this barrier has broken down and tumor cells have invaded the stroma (Figure 1A). DCIS comprises 20% of new breast cancer diagnoses, but unlike IBC, in itself is not a life-threatening disease. However, if left untreated, up to half of these patients will develop IBC within 10 years (Betsill et al., 1978; Erbas et al., 2006; Eusebi et al., 1994; Page et al., 1982; Ryser et al., 2019).

**Figure 1.**
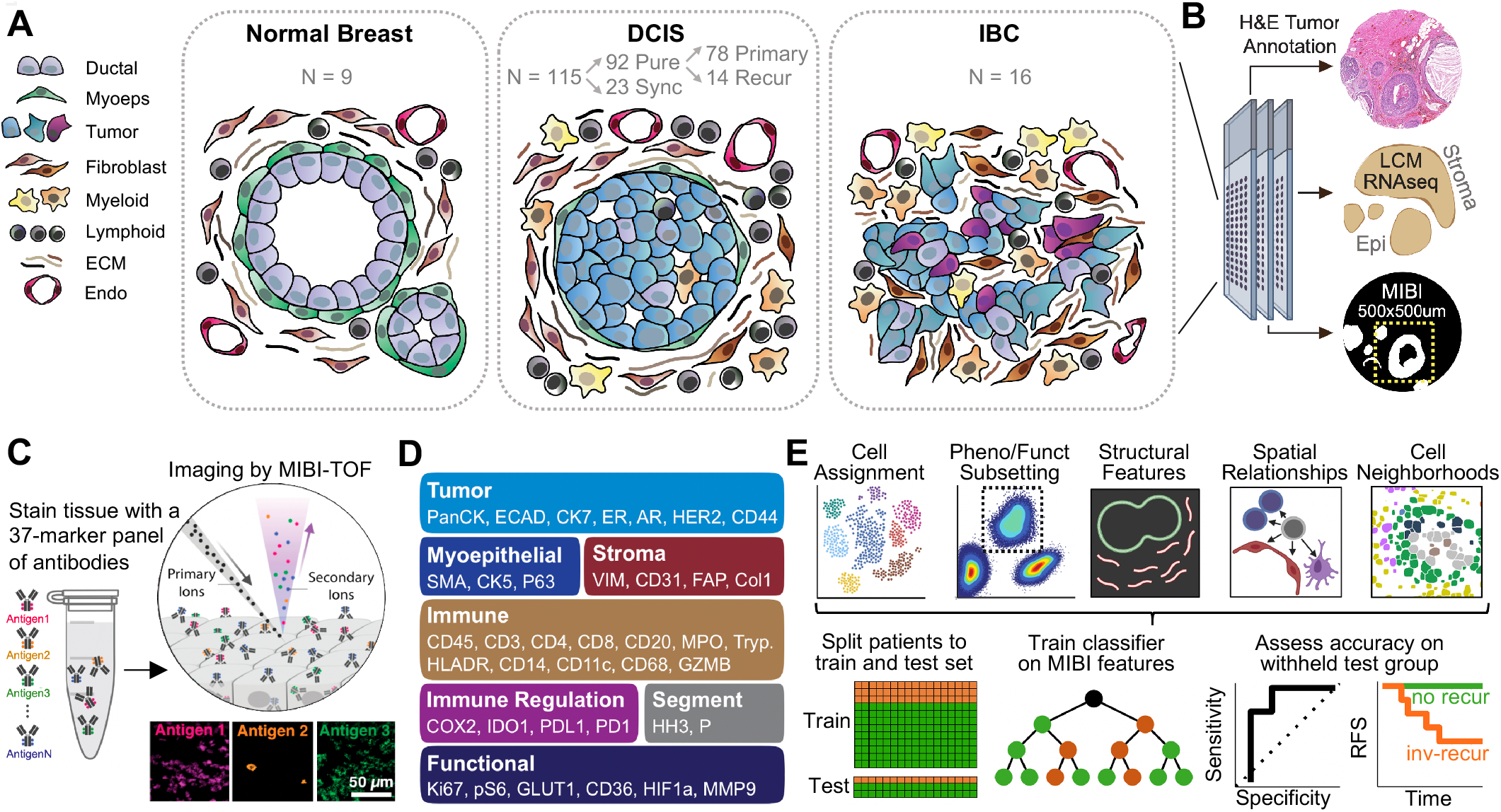
A multiplexed Imaging Interrogation of DCIS Progression to Invasive Disease. **A.** Schematic depicting the tumor stages and patient sample numbers profiled in this study, including normal breast, pure DCIS (primary or recurrent), synchronous DCIS (Sync), and invasive breast carcinoma (IBC). **B.** Depiction of the parallel tissue analysis methods used in this study including H&E, laser capture microdissection (LCM) RNAseq, and MIBI-TOF. **C.** Overview of the MIBI-TOF workflow. **D.** Markers used in the MIBI-TOF panel are displayed, grouped by target cell type or protein class. **E.** Workflow showing feature types extracted from the MIBI-TOF analysis that were used to train a random forest classifier to differentiate DCIS samples with or without risk of recurrence.

Sequencing-based approaches have been used extensively over the last decade to identify molecular features that could elucidate the connection between DCIS and IBC. Genomic profiling has identified recurrent copy number variants (CNV) that are more prevalent in high grade DCIS lesions (Afghahi et al., 2015; Buerger et al., 1999; Fujii et al., 1996). Meanwhile, comparison of paired DCIS and IBC lesions from the same patient has provided clues into the clonal evolution from *in situ* to invasive disease by revealing genomic alterations that are acquired during this transition (Ak et al., 2018; Kim et al., 2015; Newburger et al., 2013). To date, however, these findings have not been found to consistently explain this transition. Similarly, the utility of tumor phenotyping by single-plex immunohistochemical tissue staining has been limited as well.

In light of this uncertainty, clinical management has trended towards treating all patients presumptively as progressors with surgery, radiation therapy, and pharmacological interventions that carry risks for therapy-related adverse events. Consequently, this approach is likely to be overly aggressive for non-progressors. Thus, understanding the central biological features in DCIS that drive the transition to IBC is a critical unmet need.

Surprisingly, despite all the information now known about the genetic and functional state of tumor cells in DCIS, histopathology remains the only reliable way to diagnose it. Thus, DCIS is an intrinsically structured entity where the spatial orientation of tumor, myoepithelial, and stromal cells is the primary defining feature that distinguishes it from other forms of breast cancer.

To understand how DCIS structure and single cell function are interrelated, we use new tools previously developed by our lab for highly multiplexed subcellular imaging to analyze a large cohort of human archival tissue samples covering the spectrum of breast cancer progression from *in situ* to invasive disease. In previous work, we used Multiplexed Ion Beam Imaging by Time of Flight (MIBI-TOF) and a 36-plex antibody staining panel to identify rule sets governing tumor microenvironment (TME) structure in triple negative breast cancer that were highly predictive of the composition of immune infiltrates, the expression of immune checkpoint drug targets, and 10-year overall survival (Keren et al., 2018).

This effort provided a framework for how TME structure and composition could be used more generally as a surrogate readout to understand the functional response to neoplasia. With this in mind, we sought to determine to what extent similar features involving myoepithelial, stromal, and immune cells in the DCIS TME might play a pivotal role in breast cancer progression. Each of these have been implicated previously to promote local invasion (Barsky and Karlin, 2005; Ibrahim et al., 2020), metastasis (Pelon et al., 2020; Shani et al., 2020), and to correlate with clinical progression (Yang et al., 2018; Zhou et al., 2018).

Here, we report the first systematic, high dimensional analysis of breast cancer progression using the Washington University Resource Archival Human Breast Tissue (RAHBT) cohort: a clinically annotated set of archival tissue from patients diagnosed with DCIS and IBC. Because the DCIS patient population is complicated by differences in age, parity status, tumor subtype, and treatment course, a well-conceived cohort design is crucial for identifying meaningful features amidst these confounding variables. In light of this, the RAHBT cohort was composed of primary DCIS tumors from women who later progressed to invasive disease that were age and year-of-diagnosis matched with control tissue from women with DCIS that did not recur.

We used MIBI-TOF and a 37-plex antibody staining panel to comprehensively define the cellular composition and structural characteristics in 122 of these samples, which included normal breast, DCIS, and recurrent IBC samples. We applied machine learning tools for multiplexed cell segmentation and spatial analytics to enumerate 16 cell populations and to quantify how these populations are spatially distributed relative to one another. Object morphometrics and high dimensional pixel clustering were used to annotate the structure of stromal collagen and to discover new myoepithelial phenotypes that track with disease progression. These findings were corroborated by transcriptomic data acquired on coregistered tissue regions isolated by laser capture microdissection.

We systematically compared these features to understand how different phenotypic and structural properties of the DCIS TME change with progression to IBC. BC progression was typified by a reduction in myoepithelial integrity, a shift in fibroblast function towards proliferative cancer-associated states (CAFs), remodeling of collagen in the extracellular matrix (ECM), and a compositional and spatial reorganization of the immune microenvironment. We used the 1,093 features quantified in these analyses to build a random forest classifier for predicting which patients would later progress to invasive disease based exclusively on the original diagnostic biopsy. This classifier demonstrated an AUC of 0.83 and was heavily weighted for stromal features that were reliant on spatial information. Taken together, this work provides new insight into potential etiologies of DCIS progression that will guide development of future diagnostics and serve as a template for how to carry out similar analyses of preinvasive cancers.

## Results

### A multiplexed imaging interrogation of DCIS progression to invasive disease

We examined the transition from DCIS to IBC by profiling accumulative changes in the phenotype, structure, and spatial distribution of myoepithelium, tissue stroma, and immune cells in archival formalin-fixed paraffin-embedded (FFPE) patient tissue of three distinct progression groups: normal breast (n = 9), IBC (n = 16), and DCIS (n = 115). These IBC samples were disease recurrences from women with a prior diagnosis of DCIS. Of the 115 DCIS samples, 78 were RAHBT patients with a new diagnosis and no signs of IBC (pure, primary), while 14 were pure DCIS recurrences (pure, recur)(Figure 1A, Table S1). The remaining 23 patients comprised a third group of synchronous lesions procured at Stanford Hospital where both DCIS and IBC were identified in different parts of the tissue at the time of diagnosis (Sync). For this set of patients, only the *in situ* component was analyzed.

1.5 mm cores of each tumor were arranged in tissue microarrays (TMAs). Three adjacent sections were then used for 1) H&E staining and annotation by a pathologist, 2) RNA transcriptome analysis of ductal and stromal regions isolated using laser-capture microdissection (LCM-Smart-3SEQ)(Foley et al., 2019), and 3) highly multiplexed imaging by MIBI-TOF of a 500×500μm field-of-view (FOV)(Figure 1B). By ensuring that each of these analyses were spatially coregistered with one another, the proteomic and transcriptomic features revealed by MIBI-TOF and LCM-RNAseq could be directly correlated to understand the interplay between single cell composition and global transcriptional programs.

For MIBI-TOF, we constructed a 37-plex staining panel of metal-conjugated antibodies that would permit us to: 1) map the lineage and spatial location of every cell, 2) identify lineage subsets of tumor, fibroblasts, and immune cells previously implicated in BC progression, and 3) characterize the composition, integrity, and morphology of myoepithelium and collagen (Figure 1D, Table S2). The panel also included 11 functional markers for annotating proliferation, activation, hypoxic signaling, as well as markers implicated in cancer immunoregulation, including PD-L1, IDO1, COX2 and PD1 (Figure S1). The features extracted in this analysis were then used to train a random forest classifier for predicting long term outcome (Figure 1E).

### A single cell phenotypic and spatial atlas of DCIS

The workflow outlined in Figure 1 enabled high-dimensional, subcellular imaging of dozens of proteins that recapitulated the tissue architecture observed in H&E (Figure 2A). Multiplexed imaging data were processed with a low-level pipeline prior to single-cell segmentation (Figure 2B, Figure S2B)(Keren et al., 2018; McCaffrey et al., 2020; Moen et al., 2019; Valen et al., 2016), which identified on average ~924 cells in each FOV (sd = 317). To determine cell location with respect to canonical histological features, we demarcated duct, stroma, and myoepithelial regions of each image based on combinatorial marker expression (Figure 2B bottom-right). Importantly, throughout this work we will be presenting cellular data either as the frequency of a parental lineage across the entire image (e.g., macrophages as % of total immune cells) or as a cell density within a particular compartment of the image (e.g., 50 fibroblasts/mm^2^ of stroma).

**Figure 2.**
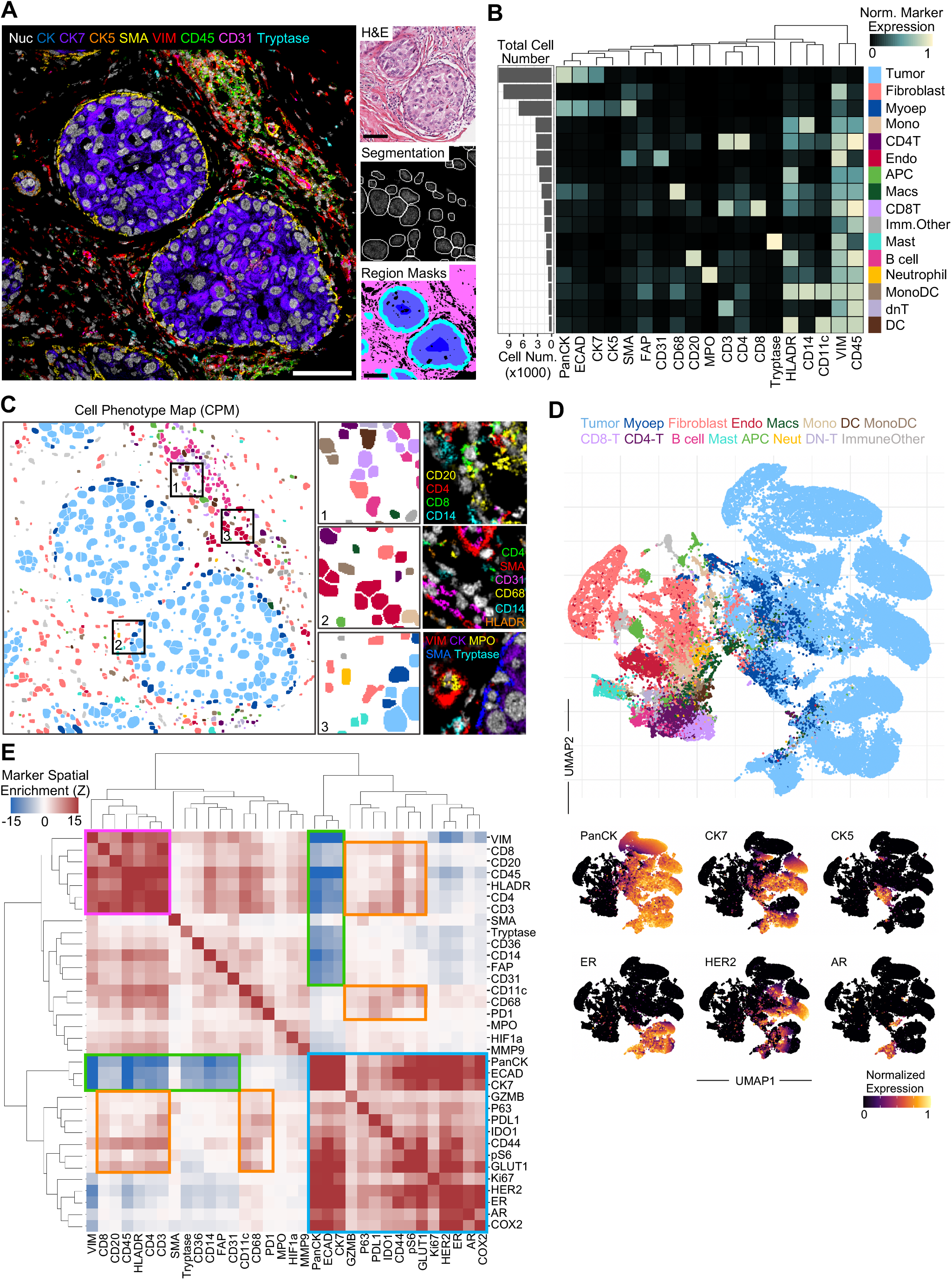
A single cell phenotypic and spatial atlas of DCIS. **A.** Representative MIBI image overlay of a DCIS tumor with a 9-marker overlay of major cell lineage markers (left) and the corresponding H&E image (top right), example of cell segmentation (middle right), and example of region masks marking stroma (pink), myoepithelial (cyan) and ductal (blue) area, scale bars = 100μm. **B.** Cell lineage assignments based on normalized expression of lineage markers (heatmap columns), rows are ordered by absolute abundance shown in the bar plot (left), while columns are hierarchically clustered (euclidean distance, average linkage). **C**. A cell phenotype map (CPM) showing cell identity by color, as defined in *F,* overlaid onto the segmentation mask. Zoomed insets with adjacent MIBI overlays show diverse lymphoid rich regions (1), endothelial-associated immune cells (2) and rare subsets like neutrophils and mast cells near ducts (3). **D.** UMAP visualization of all cell type populations in DCIS tumors (top), colored by cell type as in *F*, with additional plots overlaid with the normalized expression of tumor lineage and functional markers used to delineate tumor subsets (bottom).

Hierarchical application of the FlowSOM algorithm (Van Gassen et al., 2015) was employed to identify 16 unique cell subsets in the dataset amongst the epithelial, stromal, and immune lineages (Figure 2B, S2B). Altogether, we assigned 95% (n = 127,451 single cells) of cells to one of these subsets that in aggregate ranged in frequency from 0.7-56%. These data were used to generate cell phenotype maps (CPM) where each cell is colored according to its subset assignment. CPM images illustrated focal enrichment of lymphocytes (Figure 2C “1”), endothelial-associated immune phenotypes (Figure 2C, “2”) and sparser subsets of periductal granulocytes that included neutrophils and mast cells (Figure 2C, “3”).

Tumor cells were the most abundant cell type in DCIS samples (60% ± 20 of all cells) and were comprised of multiple subsets that were defined by variable expression of the luminal and basal lineage markers (CK7 and CK5, respectively), as well as ER, AR, and HER2 (Figure 2D). Since these cells are isolated by a layer of myoepithelium, by definition the tissue structure of DCIS is highly compartmentalized. In order to determine if our analyses were capturing this fundamental facet, we used an unbiased computational approach to identify sets of proteins that colocalize or avoid one another more frequently than would be expected by chance. Consistent with the compartmentalized nature of DCIS, tumor cell markers were spatially enriched (PanCK, ECAD, CK7, HER2, ER, AR, Figure 2E, blue box) and segregated from vascular, fibroblast, and immune markers (Figure 2E, green box). With respect to the latter, lymphoid markers demonstrated the most prominent spatial enrichment (Figure 2E, magenta box). These analyses also revealed moderate preferential enrichment in tumor positive regions for pS6, COX2, and Ki67, while immunoregulatory markers were more evenly dispersed between tumor and immune-enriched regions (Figure 2E, orange box).

### A tumor cell phenotypic switch marks invasive transition

Tumor heterogeneity in breast cancer can manifest as variations in the level of hormone receptor expression and the degree of luminal, basal, and mesenchymal differentiation. DCIS has been shown to vary across the full spectrum of both of these axes, which can confound identification of conserved features correlating with clinical outcome. In order to understand how this heterogeneity manifests in pure DCIS and throughout the transition to invasive disease, we first examined the distribution of DCIS subtypes with respect to hormone receptor status (ER, AR), HER2, and Ki67 proliferation index. These markers were robustly expressed in DCIS tumors (Figure 3A) and showed expected inter-patient variability. Using clinical cutoffs as a guide (Figure S3A), we subtyped tumors as Luminal A (ER^+^, HER2^−^, Ki67^−^), Luminal B (ER^+^, HER2^−^, Ki67^+^), HER2E (ER^−^, HER2^+^), ERHER2 (ER^+^, HER2^+^), and TNBC (ER^−^, HER2^−^) based on the frequency of positive cells for each marker. All subtypes were present in both DCIS and IBC, with similar numbers of luminal samples in each progression group (Figure 3B). HER2^+^ tumors were more predominant in DCIS, while TNBC was more prevalent in IBC (Figure S3B-C).

**Figure 3.**
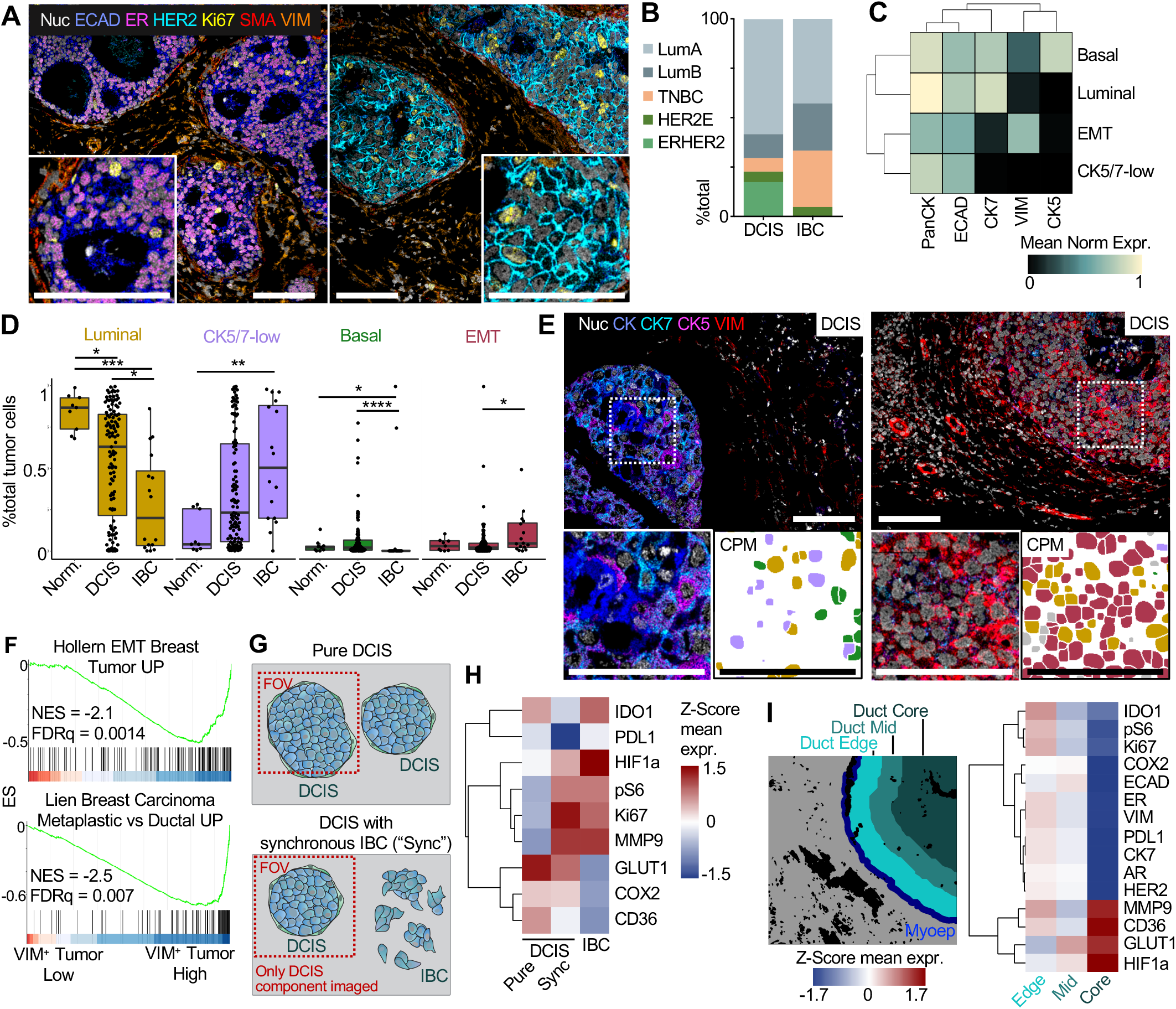
A tumor cell phenotypic switch marks invasive transition. **A.** Representative MIBI image overlays showing an ER+HER2-tumor (left) and ER-HER2+ (right), scale bars = 100μm. **B.** Stacked barplot showing the distribution of intrinsic breast cancer subtypes in DCIS and IBC tumors, as defined by receptor expression. **C**. Tumor phenotype assignments based on normalized expression of markers related to markers of tumor differentiation (heatmap columns). **D.** Frequency of tumor differentiation states across normal breast, DCIS, and IBC. **E.** Representative MIBI image overlays of DCIS tumors with basal and mesenchymal features, respectively. Zoomed insets (left) with paired cell phenotype maps (right) colored by tumor phenotype identity as in *D,* scale bars = 100μm. **F.** Geneset enrichment analysis comparing VIM-high and low tumors with genesets related to mesenchymal tumor differentiation. **G.** Schematic showing the imaging FOV location in pure and synchronous DCIS tumors, which only included the DCIS component. **H.** Heatmap of z-score normalized functional marker expression between tumor progression groups. **I.** Heatmap of z-score normalized functional marker expression in DCIS tumors comparing tumor cells on the outer duct edge, tumor cells in the duct middle (duct mid), and tumor cells in the duct core.

On comparing epithelial differentiation states in each progression group, we identified a consistent trend towards reduced luminal cell identity throughout tumor progression. Distinct phenotypic subsets of luminal (CK7^+^), basal (CK5^+^), EMT-like (VIM^+^), and CK5/7-low cells were observed in the epithelial lineage (Figure 3C). While the majority of ductal cells in normal breast were consistently luminal (84% ± 11) (Figure 3D), the composition in DCIS varied widely between being predominantly luminal or CK5/7-low (57% ± 33, 36% ± 33 respectively). In comparison to normal tissue and IBC, these lesions were also enriched with a minority fraction of basal cells (6.1% ± 11.9). With progression to IBC, CK5/7-low cells predominate more frequently and were accompanied by a relative increase in EMT-like cells that express vimentin (Figure 3E). We further examined a subset of patients with high frequencies of vimentin-positive tumor cells by LCM-RNAseq. Consistent with the shift to a mesenchymal phenotype captured by MIBI-TOF, geneset enrichment analysis (GSEA) revealed upregulation of signaling pathways relating to mesenchymal breast tumor histology and tumor invasion in patients with high vs low frequencies of VIM^+^ tumor cells (Hollern et al., 2018; Lien et al., 2007; Poola et al., 2005)(Figure 3F, Figure S3D).

The coordinated changes in tumor phenotype illustrate how cell differentiation during BC progression may follow an orderly trajectory. To further explore this possibility, we compared tumor cell functional states in pure, DCIS synchronous DCIS, and IBC. Synchronous DCIS describes lesions where distinct areas of tissue contained either fully encapsulated tumor cells (i.e., DCIS) or areas of local invasion (i.e., IBC) were both present at the time of diagnosis, but in different areas of tissue (Figure 3G). Consistent with their more aggressive behavior, DCIS tumor cells from synchronous lesions demonstrated an intermediate functional profile, with features overlapping between pure DCIS (GLUT1, CD36, COX2) and IBC (Ki67, pS6, HIF1α, MMP9) (Figure 3H).

It is not well understood how these functional states are affected by the location of tumor cells within the duct of carcinoma *in situ*, where interior tumor cells far from the duct edge may have limited access to nutrients and oxygen. Interestingly, we found almost all proliferative and cell signaling molecules to be enriched in tumor cells on the duct edge, whereas HIF1α and metabolite import receptors GLUT1 and CD36 were enriched in cells in the duct core, consistent with an adaptation to a low nutrient, hypoxic environment (Figure 3I).

### Myoepithelial breakdown and phenotypic change during DCIS progression

To understand how the structure and function of this key cellular barrier changes with progression to IBC, we next performed a targeted analysis characterizing myoepithelial cells which circumscribe both normal breast ducts and tumor cells in DCIS. Breast myoepithelium in normal tissue is a thick, highly cellular layer between the stroma and ductal cells (Figure 4A). In DCIS, the myoepithelium is notably thinned out and reduced in cellular density (Figure 4A-B). The remaining myoepithelial cells in DCIS tumors were found to have higher proliferation relative to normal tissue, with synchronous tumors having the highest levels of the Ki67 positivity of these three groups (Figure 4C).

**Figure 4.**
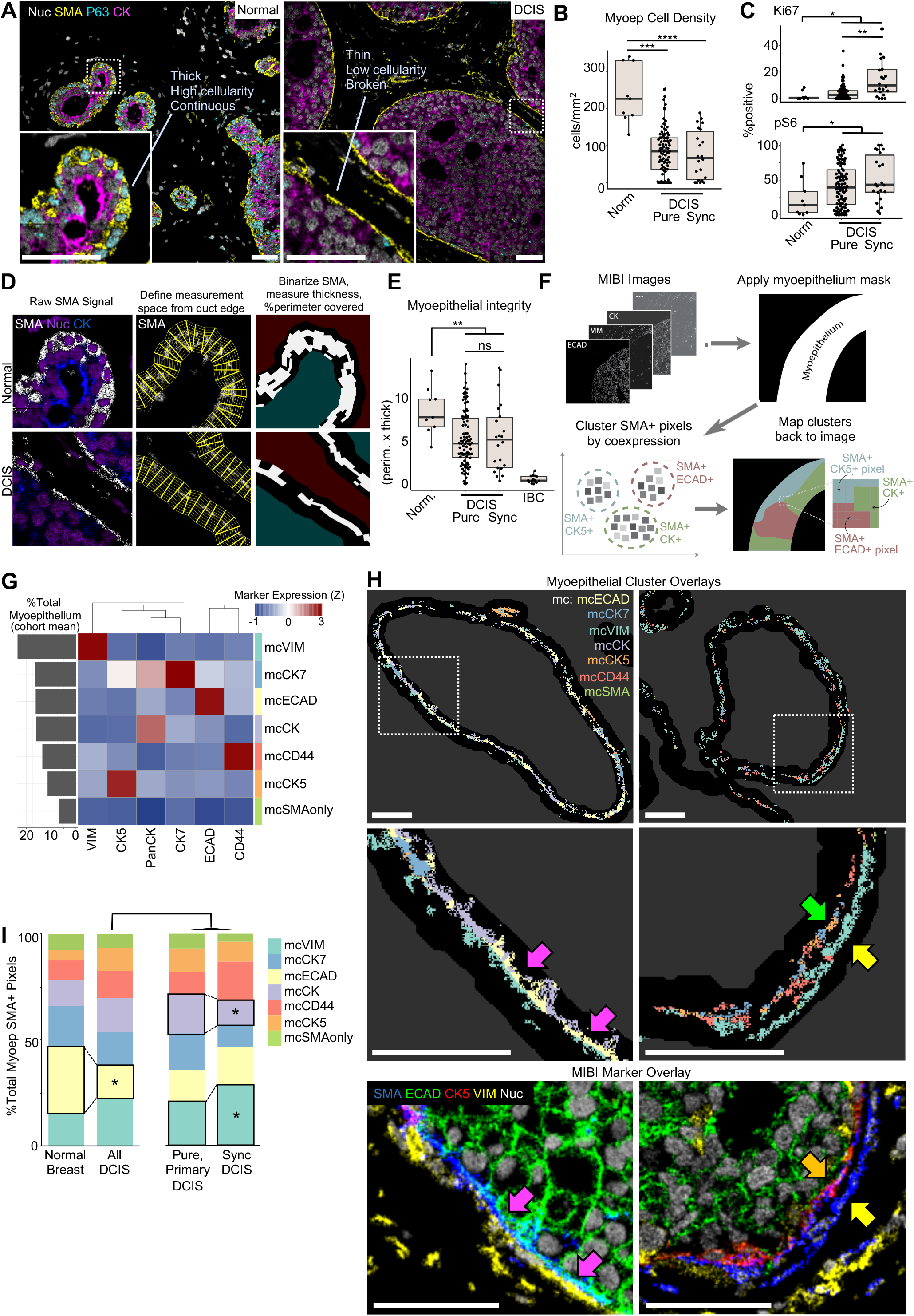
Myoepithelial breakdown and phenotypic change during DCIS progression. **A.** Representative MIBI image overlays showing SMA (yellow), p63 (cyan), and PanCK (magenta) expression in myoepithelium in normal breast (left) and DCIS (right), scale bars = 50μm. **B.** Myoepithelial cell density (cell/mm2) was quantified in periductal regions is shown for normal breast, pure DCIS, and synchronous DCIS samples. **C.** The frequency of Ki67 (top) and pS6 (bottom) positivity is compared between groups as in *B.* **D.** Illustration of workflow for quantifying myoepithelial thickness and continuity. **E.** Boxplot showing myoepithelial integrity (percent coverage × average thickness) for normal tissue and patients with pure or synchronous DCIS. **F.** Workflow schematic for pixel-based clustering of myoepithelial phenotype. **G.** Heatmap showing frequency and average marker expression for 7 myoepithelial pixel clusters (mc) with a bar plot (left) of mc abundance out of total identified myoepithelium in the cohort. **H.** *Top.* Pseudo-colored image illustrating the spatial distribution of myoepithelial pixel clusters defined in *G* for a pure (left) and synchronous (right) DCIS tumor, scale bars = 50μm. *Middle*. Magnified periductal region with mcECAD (pink arrows), mcCK5 (orange arrow), and mcVIM (yellow arrow) areas denoted. *Bottom.* Coregistered color overlays showing variations in coexpression of SMA, ECAD, CK5, and VIM corresponding to pixel cluster assignments, scale bars = 50μm. **I.** Area plots comparing the frequency of each myoep cluster across normal breast, pure, and synchronous DCIS.

Given these findings, we hypothesized that loss of myoepithelial integrity (thickness × percentage of duct-perimeter covered) in synchronous DCIS lesions would also be greater than in pure DCIS. To explore this question, we developed a new image analysis tool to quantify myoepithelial thickness and percent coverage of the duct edge (Figure 4D, see *Myoepithelial Coverage and Thickness Analysis* in Methods). This analysis revealed significant loss in myoepithelial integrity in DCIS tumors relative to normal tissue. To our surprise, however, no significant difference was observed between pure and synchronous disease. Thus, *in situ* tumorigenesis is accompanied by a reduction of myoepithelial cell density and myoepithelial integrity independent of the presence of a neighboring invasive component.

After quantifying these changes in myoepithelial structure, we next sought to determine how the function of this regulatory barrier is altered with disease progression. Due to their thin, elongated, and non-spherical cell bodies, myoepithelial cells are inherently challenging to profile with classical nuclear-based segmentation approaches which have been optimized for more conventional, ovoid cell shapes. Consequently, outlines for myoepithelial cells predicted by these methods often extend significantly beyond the true cellular border to erroneously include pixels from neighboring epithelial and stromal cells. These errors propagate in downstream cell clustering analyses to result in inaccurate phenotypic descriptions that are biased by what proteins are expressed by closely approximated neighboring cells.

To avoid this pitfall, we created a new computational approach that assigns phenotypes at the level of single pixels, rather than for whole cells (Figure 4F, see *Myoepithelial Pixel Clustering Analysis* in Methods). This strategy yielded 7 distinct, SMA^+^ myoepithelial pixel clusters (mc) defined by coexpression of PanCK, ECAD, CK7, CK5, VIM, or CD44, with SMA (Figure 4G). Mapping these pixel clusters back onto the original images revealed that multiple expressional states can exist along the perimeter of a single duct, from ECAD^+^ and CK5^+^ expression states often observed with apical preference (Figure H, pink and green arrows), and more mesenchymal states that exhibited a basal preference (e.g., VIM^+^, CD44^+^, yellow arrows). Notably, this analysis also revealed a transition from a more luminal-like state in normal samples to a more mesenchymal-like state in synchronous DCIS that aligned with analogous shifts in tumor cell differentiation and function (Figure 4I).

### Fibroblast transition and collagen architecture remodeling during DCIS tumorigenesis and progression

In light of previous studies revealing a functional and structural interdependence between myoepithelium and the surrounding stroma (Jones et al., 2003; Morsing et al., 2020), we next sought to determine if the progressive loss of myoepithelial integrity observed here correlated with changes in fibroblast function and extracellular matrix remodeling (ECM). Single cell clustering revealed four fibroblast populations that included normal (CD36 high), resting (VIM-only), myofibroblast (SMA^+^), and CAF (FAP^+^) subsets (Figure 5A). No significant differences in stromal cell density between progression groups were identified when treating fibroblasts as a single cell population (Figure 5B). However, on comparing the frequency of fibroblast subsets in normal tissue and DCIS, CAFs were found to significantly increase across tumor progression as resting fibroblasts decreased (Figure 5C), with pure DCIS tumors having a heterogeneous mixture of these two states (Figure 5D, normal fibroblasts with light blue arrows, CAFs with dark blue arrows). A corresponding increase in Ki67^+^ fibroblasts suggests that this shift in identity is driven in part by CAF proliferation (Figure 5E), which is accompanied by an increase in protein translation (high pS6). We confirmed this relationship by comparing the CAF frequency in samples with high and low pS6 and Ki67 (Figure S4A-B).

**Figure 5.**
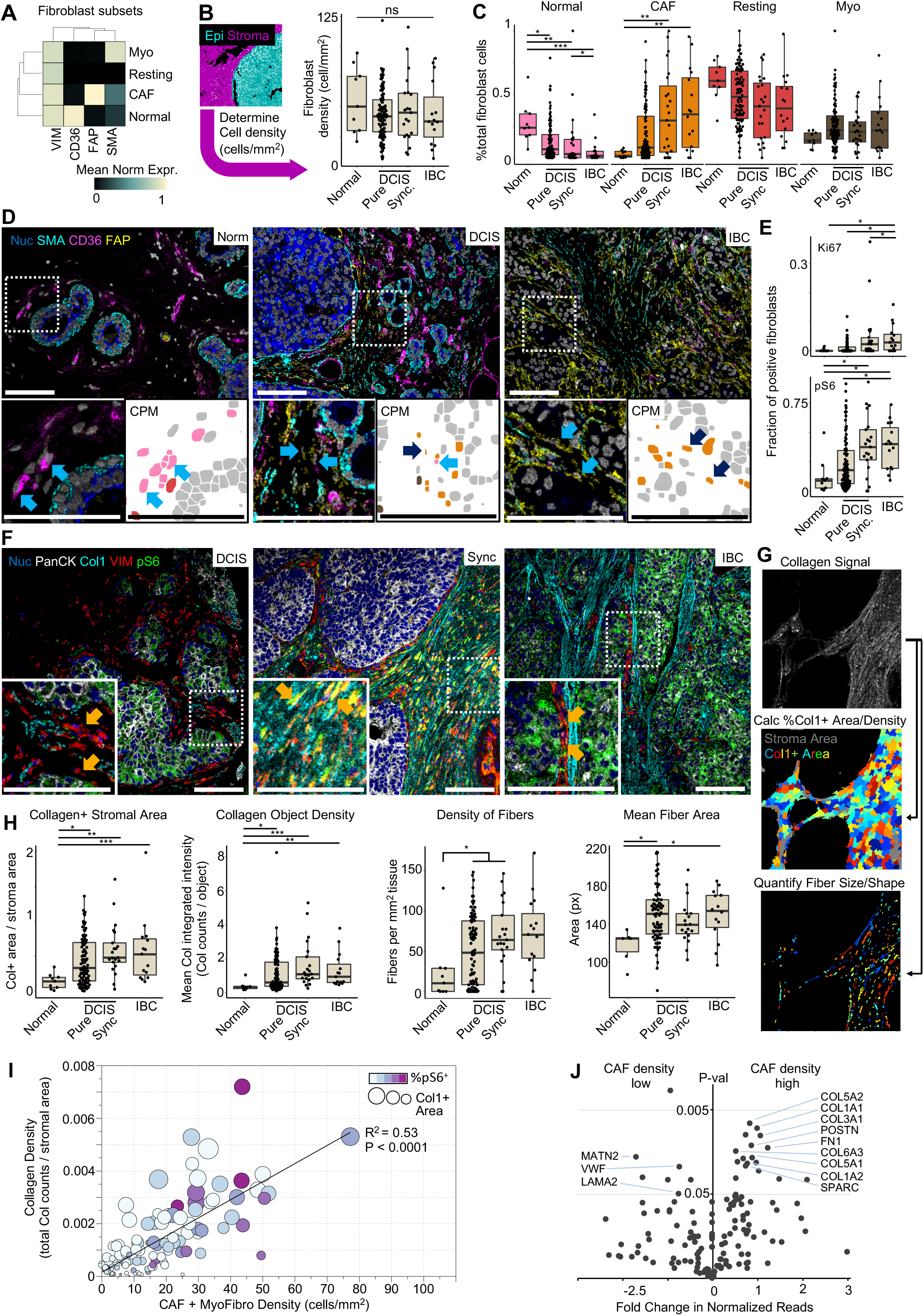
Fibroblast transition and collagen architecture remodeling during DCIS tumorigenesis and progression. **A.** Heatmap showing normalized marker expression for four fibroblast cell subsets: myofibroblasts (Myo), resting fibroblasts (Resting), cancer-associated fibroblasts (CAFs) and normal fibroblasts (Normal). **B.** *Left.* Example epithelial (cyan) and stromal (magenta) masks used to quantify stromal fibroblast density. *Right.* Boxplot of fibroblast density between tumor progression groups. **C.** Boxplots of fibroblast subset frequency across tumor progression groups. **D.** Representative MIBI image overlays showing normal, pure DCIS, and sync DCIS tumors with fibroblast markers. Zoomed insets (*left*) have paired cell phenotype maps (CPM, *right*) colored by fibroblast identity as in *C,* scale bars = 100μm. **E.** The frequency of Ki67 and pS6 positivity in fibroblasts is shown across progression groups. **F.** Representative MIBI image overlays showing VIM+ fibroblasts (red) with varying levels of pS6 expression (green) and nearby collagen 1 (Col1, cyan) deposition, scale bars = 100μm. **G.** Schematic showing the quantitation of MIBI collagen signal to identify %collagen+ stromal area, collagen density, and collagen fiber morphometrics. **H.** Collagen+ stromal area, collagen density, collagen fiber density (fibers/mm2) and fiber area are quantified across tumor progression groups. **I.** Scatterplot comparing summed density of CAFs and myofibroblasts versus collagen density. Size and color of points are proportional to collagenized area and fibroblast pS6 positivity, respectively. **J.** Volcano plot of ECM-related gene expression for the top and bottom CAF-enriched DCIS tumors.

Given these findings, and that dense fibrillar collagen often appeared to be juxtaposed with pS6^+^ fibroblasts in progressed tumors (Figure 5F, orange arrows), we next sought to determine how collagen remodeling was related to CAF location, frequency, and phenotype. To achieve this, we developed new computational tools for collagen morphometrics that were used to determine the shape, length, and density of individual fibers (Figure 5G, see *Collagen Morphometrics* in Methods). These analyses revealed that DCIS and IBC tumors had higher collagen density and longer fiber length compared to normal breast (Figure 5H), suggesting that collagen deposition and fibrillar remodeling were coordinated with the phenotypic shift to CAFs. Indeed, direct comparison of collagen density and collagen-positive area to the density of CAFs and myofibroblasts in the stroma revealed a strong correlation (Figure 5I). Furthermore, pS6^+^ fibroblasts were also enriched in these collagen and CAF-dense tumors. Together these data suggest a direct relationship between CAF activation and collagen deposition and remodeling.

Finally, to identify which specific collagen isoforms correlate with this activity and to determine if additional ECM proteins are involved, we compared ECM transcript levels in stroma of CAF-high- and low-density tumors using LCM RNAseq. We found the majority of collagen species were upregulated in CAF-high tumors with COL5A2 and COL1A1 being the most significant of these, consistent with MIBI-TOF quantitation of COL1A1 protein (Figure 5J). In addition, CAF-dense tumors showed increased deposition of fibronectin (FN1), SPARC and periostin (POSTN), indicative of CAF-remodeling and a shift towards a pro-invasive stroma (Barth et al., 2005; Malanchi et al., 2012).

### Characterizing the preinvasive immune microenvironment and its compartmental evolution throughout progression

Having identified coordinated shifts in tumor differentiation, myoepithelial integrity, and fibroblast function, we next sought to understand how immune composition changed with disease progression. We found monocytes, mast cells, and HLA-DR^+^ antigen presenting cells (APCs) to be the most abundant immune cells in pure DCIS (Figure 6A). Immune cells were typically found in the stroma and were occasionally embedded in ducts (Figure B, orange arrow). To quantify the spatial distribution of immune cells in these compartments, we interrogated cell density in epithelial and stromal mask regions (Figure 6C). This analysis identified a clear stromal preference when treating immune cells as a single population (Figure 6D, S5A). To understand if this preference remained valid when considering specific subsets of lymphoid and myeloid cells, we compared the local frequency within stromal and ductal regions for each cell type. CD4^+^ T cells, B cells, monocytes, APCs and mast cells all demonstrated a statistically significant stromal preference, while macrophages were significantly enriched in ductal regions (Figure 6E). Interestingly, differential enrichment of CD4^+^ and CD8^+^ T cells resulted in a CD4/CD8 ratio that skewed towards CD8^+^ T cells in ducts and CD4^+^ T cells in stroma (Figure 6F).

**Figure 6.**
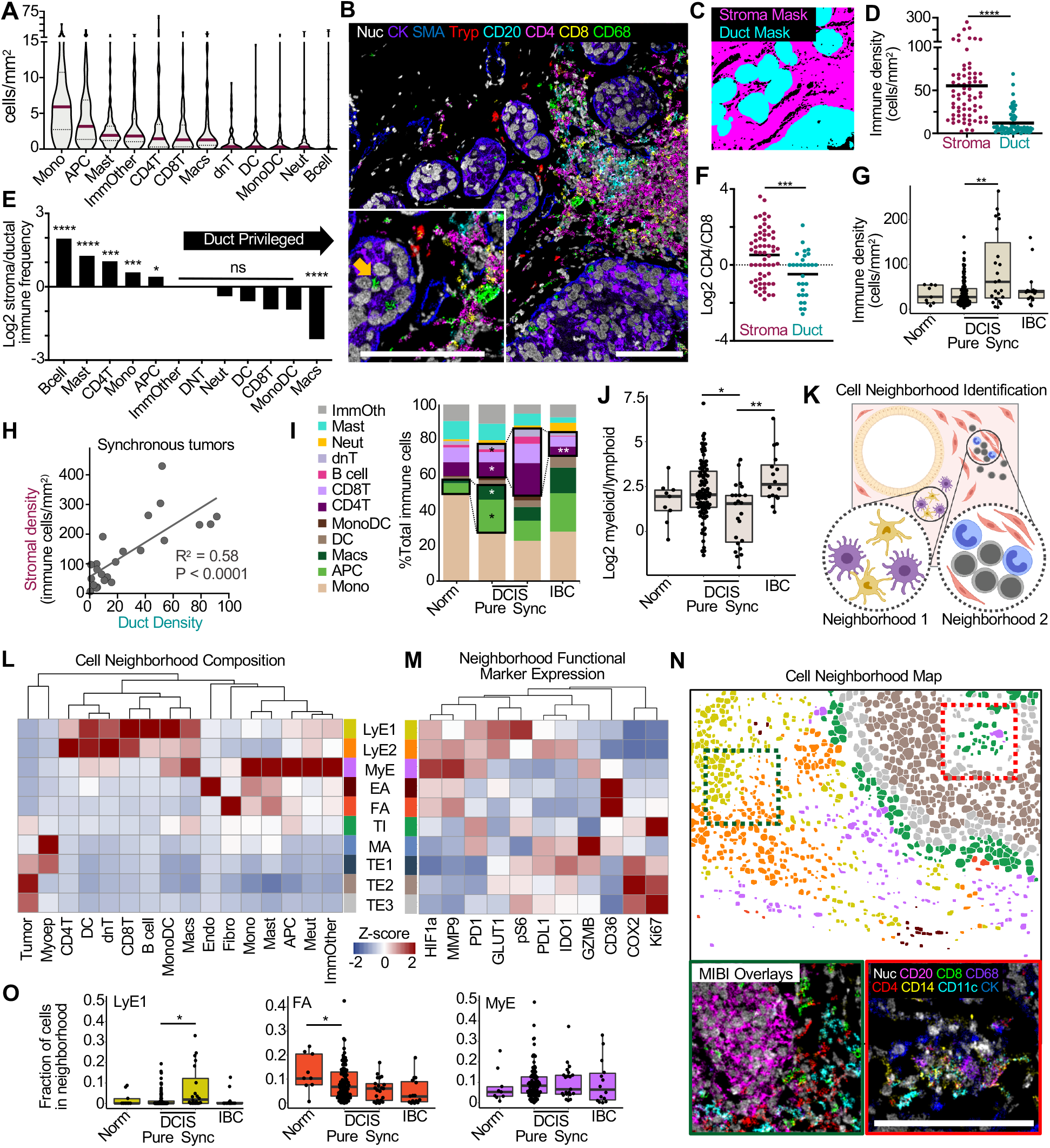
Characterizing the preinvasive immune microenvironment and its compartmental evolution throughout progression. **A**. Violin plot examining immune cell density in pure DCIS, ranked by median density per patient. **B**. Representative MIBI image overlay of a pure DCIS tumor with major immune cell type markers, inset and arrow highlighting intraductal immune phenotypes. **C.** Mask overlay showing delineation of stroma and duct regions in *B,* scale bars = 100μm. **D.** Scatterplot comparing immune cell density between the stroma and duct compartments per patient. **E.** Column plot showing the ratio (Log2) of immune cell type frequency between stroma and ductal compartments, ranked from high (stromal preference) to low (duct preference). Asterisks denote significance comparing compartment frequency of a given cell type across all pure DCIS patients. **F.** Log2 ratio of CD4+ to CD8+ T cells is displayed per patient for the stroma and duct compartments. **G.** Whole image immune density is compared across tumor progression groups. **H.** Scatterplot comparing stromal and ductal immune density per patient in synchronous tumors. **I.** Area plot showing the change in immune subset frequency across progression groups. Effector-myeloid cell subsets are boxed and compared between normal breast and pure DCIS tumors; asterisks denote significant differences in frequency. Lymphocyte subsets are boxed and compared between pure DCIS, synchronous DCIS, and IBC, asterisks denote significance vs the synchronous group. **J.** Boxplots showing the log2 ratio of myeloid to lymphoid cells in tumor progression groups. **K.** Illustration depicting different spatially-enriched cellular neighborhoods. **L.** Heatmap showing z-score normalized cell type frequency for each cellular neighborhood: lymphocyte-enriched (LyE1, LyE2), myeloid-enriched (MyE), endothelial-associated (EA), fibroblast-associated (FA), tumor-interface (TI), myoepithelial-associated (MA), and tumor-enriched (TE1-3). **M.** Heatmaps showing z-score normalized mean expression for functional markers in each cellular neighborhood. **N.** *Top.* Cell neighborhood map showing the spatial localization of distinct neighborhoods, denoted by color as in *M. Bottom.* Color overlays for lymphocyte-enriched (green dotted line) or tumor-interface (red dotted line), scale bar = 100μm. **O.** Boxplot showing frequency of cells assigned to LyE1 (yellow), Fibroblast-associated (Red) and MyE (purple) cell neighborhoods across tumor progression groups.

We next investigated how immune cell prevalence and spatial enrichment evolves with transition from *in situ* tumorigenesis to invasive disease by comparing pure DCIS with synchronous lesions and IBC. Immune cell density was significantly increased in synchronous lesions compared to all other groups (Figure 6G). Notably, this increase in immune infiltrate was present in both the stroma and ducts of these lesions (Figure 6H), suggesting a coordinated influx into the ducts during increased stromal immune infiltration. By comparing the cell density for each immune cell subset with respect to disease stage, we observed an increase in effector myeloid cells (Macs, APC) in pure DCIS compared to normal breast (Figure 6I). Importantly, this also revealed the increase in immune infiltrate in synchronous tumors to be driven primarily by an influx of B and T lymphocytes (Figure 6I, S5B), resulting in an immune microenvironment more skewed towards lymphocytes (Figure 6J). Subsequently, both T cell frequency and myeloid to lymphoid ratio in IBC tumors return to values similar to pure DCIS.

In order to better understand how this feature and other immune programs were spatially organized, we applied a K-means clustering approach to identify distinct cellular neighborhoods (CNs), where a CN is defined by a set of cell types found to spatially co-occur across the cohort (Figure 6K, see *Protein and Cellular Spatial Enrichment Analyses* in Methods). Through this approach, we identified 10 CNs that we categorized as being lymphocyte-enriched (LyE1, LyE2), myeloid-enriched (MyE), endothelial-associated (EA), fibroblast-associated (FA), myoepithelial-associated (MA), tumor-interface (TI), and tumor-enriched (TE1-3, Figure 6L-N).

Interestingly, single cell expression of functional markers was found to be correlated with CN, even though these parameters were not included in the K-means neighborhood assignment analysis. For example, HIF1α and MMP9 expressing cells were enriched in MyE, while the frequency of pS6^+^ cells was highest in LyE1 (Figure 6L). Macrophages were a constituent of numerous CNs and showed functional state distinction based on neighborhood association, including increased PDL1 expression within LyE neighborhoods, in addition to pS6 (Figure S5C-D). Notably, the LyE1 neighborhood was also enriched for T and B cells, consistent with tertiary lymphoid structure formation (see Figure 6N, bottom left). In line with the trends observed for T cell infiltrates, we found the frequency of cells belonging to LyE1 to be increased in synchronous lesions (Figure 6O). Taken together, these findings indicate that early stromal invasion in synchronous tumors triggers an influx of T cells and formation of TLS structures. We find that by IBC, however, the tumor immune microenvironment has reverted to a myeloid-skewed, immunosuppressed state with diminished T cell presence.

### Identifying DCIS features correlated with recurrence outcomes

Having extensively quantified the multi-compartmental cellular and structural elements of DCIS tumors, we leveraged these data to identify features associated with the risk of recurrence following primary DCIS resection. We selectively examined these features in diagnostic tissue procured at the time of initial presentation in two sets of patients. The first set, referred to as “case”, consisted of 31 patients who had a recurrence (DCIS or IBC) within 2-15 years of being treated for newly diagnosed pure DCIS. The second set, referred to as “control”, consisted of 47 patients with pure DCIS that did not recur within 11+ years.

Using these outcome groups and 1,093 phenotypic, functional, spatial, and morphologic features extracted from our MIBI-TOF analyses (Table S3), we trained two random forest classifier models. The first was an all-recurrence model for predicting which patients would have a recurrence of DCIS *or* IBC. The second was an invasive recurrence model for predicting IBC recurrence *exclusively* (Figure 7A). Low observation and overly correlated features were removed from the dataset and the patient population was randomly split 80/20 to training and test groups. We evaluated classifier accuracy in the withheld test set, where the all-recurrence and invasive models achieved an AUC of 0.79 (CI 0.51:1) and 0.83 (CI 0.59:1), respectively (Figure 7B). When stratifying patients by their predicted labels, we found a significant difference in recurrence probability over time (Fig. 7C, Figure S6A), with no recurrence events in the patients predicted by the invasive model to be non-progressors. Although sample size precluded us from being able to eliminate patient demographics and differences in clinical therapy as a confounder in this analysis, treatment regimens known to affect recurrence rates (i.e., mastectomy, radiation, tamoxifen) were well distributed between the case and control patients (Figure S6B). Likewise, no significant difference in classifier predictions were identified with respect to these variables (Figure S6C).

**Figure 7.**
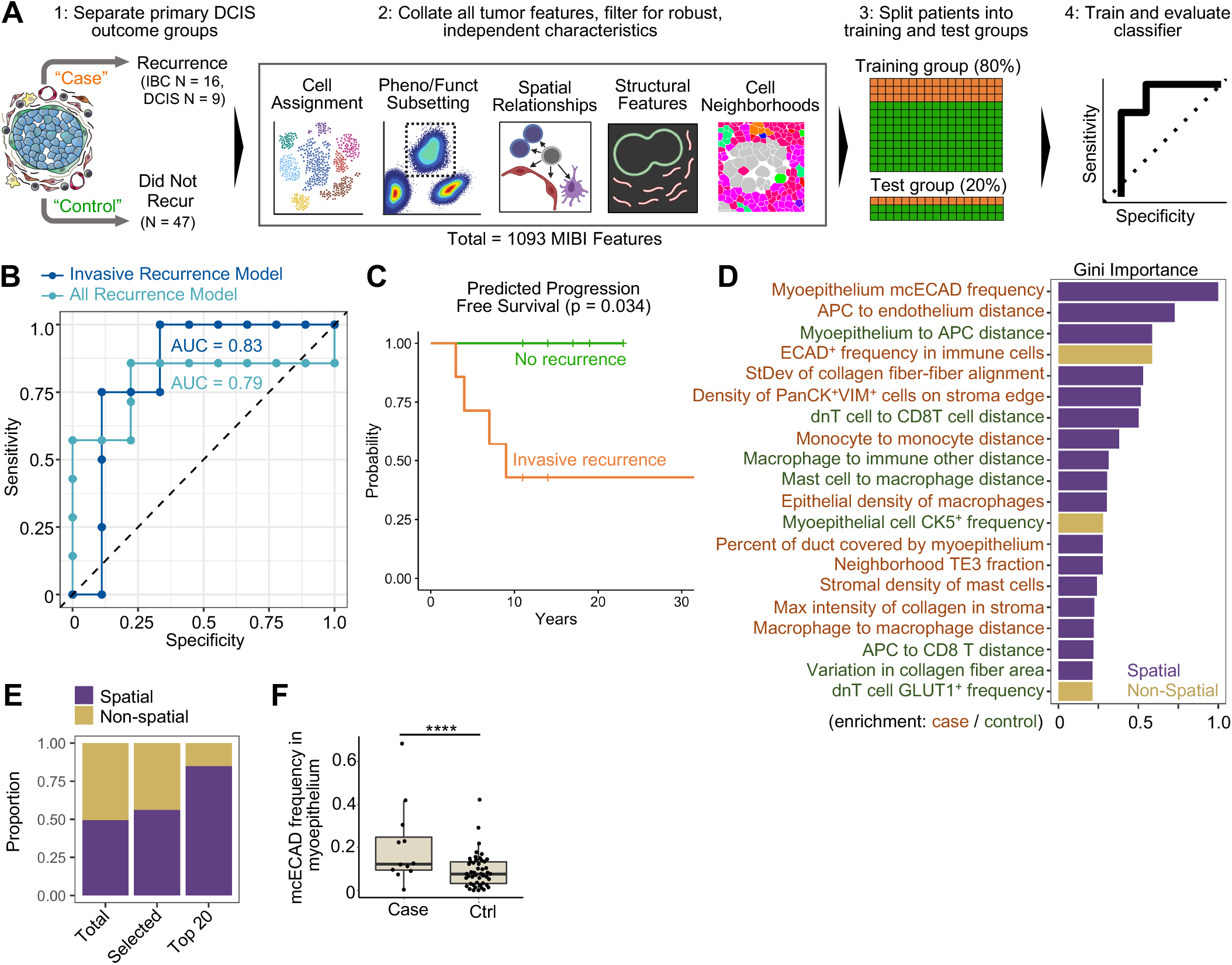
Identifying DCIS features correlated with recurrence outcomes. **A.** Schematic illustrating the different outcome groups of primary DCIS including “cases” that recurred either as IBC or DCIS, and “controls” with no recurrence in >11yr follow-up. 1,093 MIBI features of numerous tumor metrics were used to train a random forest classifier to differentiate case and control samples. Classifier specificity was then tested on a withheld 20% of patients. **B.** AUC plot showing classifier sensitivity and specificity. **C.** Predicted survival of patients identified in the test set of the invasive-recurrence model as case or control. **D.** MIBI features with top classifier importance for the IBC recurrence model are shown, ranked by Gini importance. Features are colored based on enrichment either in cases (orange) or controls (green), importance bars are colored based on the feature utilizing spatial information (purple) or not (gold). **E.** The distribution of spatial vs non-spatial features are shown for all features identified (total), those used by the model (selected), and those in the top 20 most important features (top 20). **F.** Boxplot showing the frequency of the mcECAD myoepithelial phenotype between invasive cases and controls.

To understand the biology being leveraged by this classifier to accurately discriminate pre-invasive from indolent DCIS tumors, we ranked the top 20 features based on Gini importance. These features primarily consisted of metrics related to the phenotype of myoepithelium, the structure of collagen fibers in the extracellular matrix, and the spatial distribution of multiple immune cell subsets (Figure 7C). Notably, spatial metrics describing cell densities, cell neighborhoods, pairwise cell distances, collagen structure, and multiplexed subcellular features were overrepresented and accounted for 17 of the top 20 metrics in the invasive model (Figure 7D, Table S3). Immune cell metrics comprised about half of these and were myeloid skewed (Figure S6D, with 9 relating specifically to myeloid subsets and 3 to lymphoid subsets. Similarly, enrichment for spatial metrics related to myoepithelium, collagen, and myeloid cells were observed in the all-recurrence model as well (Figure S6D-F). Stromal density of PanCK^+^VIM^+^ cells ranked in the top 20 features. These cells were rare (median of 0 in case and controls) and on manual inspection appeared to represent fibroblasts where PanCK expression from closely neighboring epithelial cells was misassigned. Interestingly, both models identified pixel-level, ECAD^+^ myoepithelial expression as the most predictive metric (mcECAD, see Figure 4). When comparing case and control samples, we found the frequency of this feature to be significantly different between these outcome groups, independent of the classifier model, and to be readily identifiable on targeted inspection of the original imaging data (p < 0.001, Figure 7F).

## Discussion

Here, we report the first multicompartmental atlas of the single cell composition and structure of DCIS. The central focus of this study was to characterize the changes undergone with progression to IBC where tumor cells breach the duct to invade the surrounding stroma. Previous work examining BC progression have attempted to attribute this transition either to tumor-intrinsic factors or to specific features of stromal cells in the surrounding TME. By simultaneously mapping both tumor and stromal cell identity and function in intact human tissue, we sought to treat the DCIS TME as a single ecosystem where progression to invasive disease depends on the spatial distribution and function of multiple cell types, rather than on any single cell subset.

Meeting this goal required first assembling a large, well-annotated, and diversified pool of human DCIS tissue: the RAHBT cohort. This effort was motivated in part by the success of similar work investigating invasive disease (i.e. METABRIC) that have provided deep insights into breast tumor composition and have served as authoritative resources in breast cancer research (Curtis et al., 2012). To achieve this, the Breast PreCancer Atlas constructed a unique set of archival human surgical resections that captured the full spectrum of breast cancer progression, from normal tissue, to pure DCIS and IBC. Assembling all of these cases into TMAs has enabled a one-of-a-kind workflow for multiomics analyses where genomic, transcriptomic, and proteomic techniques are performed not only on the same samples, but on coregistered serial sections of the same local region of tissue.

Here, we describe the first major analysis of the RAHBT cohort where high dimensional imaging was used to characterize BC progression. We used MIBI-TOF for subcellular imaging of 140 tumor and normal breast samples using a 37-marker staining panel (122 and 23 samples from RAHBT and Stanford cohorts, respectively). Tumor cell differentiation and function were found to transition along a continuum from pronounced luminal features in normal breast to a more undifferentiated, cytokeratin-low state in invasive disease that had increased mesenchymal features. This shift was accompanied by an upregulation of HIF1α, MMP9, and IDO in tumor cells, which have been shown to directly elicit EMT, promote invasion, and drive immune tolerance, respectively (Kolijn et al., 2018; Lin et al., 2011; Peng et al., 2018; Zhang et al., 2015, 2019). With transition to DCIS, the frequency of an E-cadherin-high myoepithelial phenotype that predominated normal breast tissue decreased, as a more mesenchymal, CD44- and VIM-high state increased. Interestingly, no difference in myoepithelial cell density or structural integrity was found when comparing DCIS in pure and synchronous lesions. Given that the invasive and *in situ* components of synchronous tumors are closely related on a genomic level (Ak et al., 2018; Kim et al., 2015; Newburger et al., 2013) these findings suggest that transition to invasion disease is regulated at least in part by the local microenvironment.

These epithelial changes were accompanied by a stromal transition towards higher numbers of activated, proliferating CAFs and densely aligned fibrillar collagen (Conklin et al., 2011; Esbona et al., 2018). Although the total immune density was comparable to normal breast tissue, DCIS tumors exhibited a shift from a monocyte-predominant environment to one enriched for APCs and intraductal macrophages. In line with recent findings by other groups (Alcazar et al., 2017; Kim et al., 2020) synchronous DCIS/IBC tumors were marked by a stromal spike in T and B cells and formation of tertiary lymphoid structures. This feature distinguishes them from the myeloid-skewed IBC samples profiled in this study. Taken together, these findings support a model for breast cancer progression where invasive disease occurs through multiple coordinated, dynamic interactions of the surrounding stroma, myoepithelium, and tumor.

Given the urgent need to better stratify DCIS patients based on risk of progression, we tested to see if these spatial and phenotypic features could be used to predict IBC recurrence based exclusively on diagnostic DCIS tissue. Using 1,093 features, we trained a random forest classifier model for identifying patients that would later progress to IBC that achieved an AUC of 0.83 on withheld test samples. Although the performance was impressive, certain caveats should be taken into account when considering how generalized this model might be. Given the complexity of breast cancer subtypes and the impact of patient demographics on outcome (Alaeikhanehshir et al., 2020; Liu et al., 2019), the sample size in this study may not have been sufficient to fully account for the confounding effects of these variables. Lastly, since all patients in the RAHBT cohort received one or more therapeutic interventions, the features leveraged by this model to identify non-progressors might not be valid when applied to patient populations where therapy is omitted.

With these considerations in mind however, these results do offer three compelling and overarching insights. First, spatial metrics relating phenotype to structure and morphology were significantly over-represented relative to non-spatial metrics, accounting for almost 85% of the top 20 features identified by the classifier model. Second, the most influential features were primarily related to the stroma rather than the tumor cells themselves. This included a previously unreported E-cadherin high myoepithelial phenotype as well as collagen fiber size and alignment with respect to the duct. Third, high ranking immune features more often related to myeloid than to lymphoid subsets, particularly those in close proximity with myoepithelium or residing inside the duct. This skewing underscores the need to better understand how macrophages promote TME immune suppression, tumor proliferation, and local invasion (Esbona et al., 2018; Goswami et al., 2005; Linde et al., 2018; Ruffell et al., 2012).

Taken together, this study offers a comprehensive, multi-compartmental atlas of preinvasive breast cancer that illustrates the full continuum of tissue structure and function starting from a homeostatic state in normal breast through *in situ* and invasive disease. Combining this comprehensive data set with extensive patient follow-up has enabled identification of tumor features that are associated with DCIS recurrence and offers a framework for exciting follow-on efforts. With this in mind, we are actively planning a larger study that will further evaluate the biological significance of spatial features relating to myoepithelium, collagen, and myeloid cells and to determine if they can be used to prospectively risk stratify patients with a new DCIS diagnosis.

## Methods

### Patient Cohort

We utilized a retrospective study cohort of patients from the Washington University Resource of Archival Tissue (RAHBT) that contained two outcome groups: controls (“Ctrl”) composed of patients with DCIS who had no recurrence and cases (“Case”) composed of patients with DCIS who had either a DCIS or an IBC recurrence. For each case, we matched two controls who remained free from recurrent lesions, based on age at diagnosis (+/− 5 years), and type of definitive surgery (mastectomy or lumpectomy). For each DCIS diagnosis we retrieved primary and recurrent tumor slides and blocks for pathology review, secured a whole slide image of each sample, marked for TMA cores, and generated TMA blocks with 84 1.5mm cores, including additional tonsil and normal breast controls.

Supplemental table 1 summarizes the data for the cases in the cohort. Median age at diagnosis was 54, year of diagnosis was 1986 to 2017, and time to recurrence with was 8.8 years for invasive lesions, and 5.3 years for premalignant lesions. For women in the cohort with no recurrence, follow up extended to 132 months, on average. Treatment of initial DCIS ranged from lumpectomy with radiation (approximately half of cases), and lumpectomy with no radiation (20%) and mastectomy with no radiation for 30%. The RAHBT cohort is composed of African American women (26%) and white women (74%). We also profiled a supplemental cohort of patients from the Stanford Hospital with synchronous (“Sync”) DCIS and IBC tumors from 2007-2009. A 216-core TMA block was generated with 1mm tumor cores, with additional tissue controls.

5μm serial sections of each TMA slide were cut onto glass slides for hematoxylin and eosin (H&E) staining, onto laser-capture slides for LCM-RNAseq (SMART-3SEQ) and cut onto gold- and tantalum-sputtered slides for MIBI-TOF imaging. H&E slides were inspected by a breast cancer pathologist to address DCIS purity and demarcate regions of DCIS to guide MIBI imaging and laser dissection of epithelial and stromal area. The Stanford Hospital cohort was without paired LCM-RNAseq analysis.

### Antibody Preparation

Antibodies were conjugated to isotopic metal reporters as described previously (Keren et al., 2018; McCaffrey et al., 2020). Following conjugation antibodies were diluted in Candor PBS Antibody Stabilization solution (Candor Bioscience). Antibodies were either stored at 4°C or lyophilized in 100 mM D-(+)-Trehalose dehydrate (Sigma Aldrich) with ultrapure distilled H2O for storage at −20°C. Prior to staining, lyophilized antibodies were reconstituted in a buffer of Tris (Thermo Fisher Scientific), sodium azide (Sigma Aldrich), ultrapure water (Thermo Fisher Scientific), and antibody stabilizer (Candor Bioscience) to a concentration of 0.05 mg/mL. Some metal-conjugated antibodies in this study were used as secondary antibodies, targeting hapten groups on hapten-conjugated primary antibodies, this included the pairs PDL1-Biotin and Anti-Biotin^149Sm^, and ER-Alexa488 and Anti-Alexa488^142Nd^. Information on the antibodies, metal reporters, and staining concentrations is located in Table S2.

### Tissue Staining

Tissues were sectioned (5μm section thickness) from tissue blocks on gold and tantalum-sputtered microscope slides. Slides were baked at 70°C overnight followed by deparaffinization and rehydration with washes in xylene (3x), 100% ethanol (2x), 95% ethanol (2x), 80% ethanol (1x), 70% ethanol (1x), and ddH2O with a Leica ST4020 Linear Stainer (Leica Biosystems). Tissues next underwent antigen retrieval by submerging sides in 3-in-1 Target Retrieval Solution (pH 9, DAKO Agilent) and incubating at 97°C for 40 minutes in a Lab Vision PT Module (Thermo Fisher Scientific). After cooling to room temperature slides were washed in 1x PBS IHC Washer Buffer with Tween 20 (Cell Marque) with 0.1% (w/v) bovine serum albumin (Thermo Fisher). Next, all tissues underwent two rounds of blocking, the first to block endogenous biotin and avidin with an Avidin/Biotin Blocking Kit (Biolegend). Tissues were then washed with wash buffer and blocked for 1 hour at room temperature with 1x TBS IHC Wash Buffer with Tween 20 with 3% (v/v) normal donkey serum (Sigma-Aldrich), 0.1% (v/v) cold fish skin gelatin (Sigma Aldrich), 0.1% (v/v) Triton X-100, and 0.05% (v/v) Sodium Azide. The first antibody cocktail was prepared in 1x TBS IHC Wash Buffer with Tween 20 with 3% (v/v) normal donkey serum (Sigma-Aldrich) and filtered through a 0.1μm centrifugal filter (Millipore) prior to incubation with tissue overnight at 4°C in a humidity chamber. Following the overnight incubation slides were washed twice for 5 minutes in wash buffer. The second day antibody cocktail was prepared as described and incubated with the tissues for 1 hour at 4°C in a humidity chamber. Following staining, slides were washed twice for 5 minutes in wash buffer and fixed in a solution of 2% glutaraldehyde (Electron Microscopy Sciences) solution in low-barium PBS for 5 minutes. Slides were washed in PBS (1x), 0.1 M Tris at pH 8.5 (3x), ddH2O (2x), and then dehydrated by washing in 70% ethanol (1x), 80% ethanol (1x), 95% ethanol (2x), and 100% ethanol (2x). Slides were dried under vacuum prior to imaging.

### MIBI-TOF Imaging

Imaging was performed using a MIBI-TOF instrument with a Hyperion ion source. Xe^+^ primary ions were used to sequentially sputter pixels for a given FOV. The following imaging parameters were used: Acquisition setting: 80 kHz, Field size: 500 μm^2^, 1024 × 1024 pixels, dwell time: 5ms, median gun current on tissue: 1.45nA Xe^+^, ion dose: 4.23 nAmp hours / mm^2^ for 500 μm^2^ FOVs.

### Low-level Image Processing and Single Cell Segmentation

Multiplexed image sets were extracted, slide background-subtracted, denoised, and aggregate filtered as previously described (Keren et al., 2018; McCaffrey et al., 2020). Nuclear segmentation was performed using an adapted version of the DeepCell CNN architecture (McCaffrey et al., 2020; Valen et al., 2016). To more effectively capture the range of cell shapes and morphologies present in DCIS, we generated two distinct segmentations for each image. The first used a radial expansion of three pixels and a stringent threshold for splitting cells (See Figure S2A, *Stroma Parameters*). The second used a radial expansion of one pixel and lenient threshold for splitting cells (*Epithelial Parameters*). We combined these masks together using a post-processing step which gave preference to the epithelial segmentation mask, overriding and stromal-mask-detected objects in the same area. Smaller cells identified by the stromal settings and missed in the epithelial settings were combined to the final cell mask. A cell nuclei (“Nuc”) channel combining HH3 and endogenous phosphorous (P) signal was made to increase signal robustness for nuclei detection.

### Single Cell Phenotyping and Composition

Single cell data was extracted for all cell objects and area normalized. Single cell data was linearly scaled by average cell area across the cohort and asinh-transformed with a co-factor of 5. All mass channels were scaled to 99.9th percentile. In order to assign each cell to a lineage, the FlowSOM clustering algorithm was used in iterative rounds with the Bioconductor “FlowSOM” package in R (Van Gassen et al., 2015). The first clustering round separated cells into 100 clusters that were subsequently merged into one of five major cell lineages (tumor, myoepithelial, fibroblast, endothelial, immune) based on the clustering nodes. Proper lineage assignments were ensured by overlaying Flowsom cluster identity with lineage-specific markers. Supervised lineage reassignment was performed where needed. Immune cells were subclustered again to delineate B cells, CD4^+^ T cells, CD8^+^ T cells, monocytes, MonoDC cells, DC cells, macrophages, neutrophils, mast cells, double-negative CD4^−^CD8^−^T cells (dnT cells), and HLADR^+^ APC cells. CD45^+^-only immune cells were annotated as ‘immune other.’ Tumor and fibroblast cells were similarly clustered again to reveal phenotypic subsets, as shown in Figure S2. Altogether, we assigned 94% (n = 127,451 of 134,631) of cells to 16 subsets, with the remaining nucleated cells with absent or very low levels of lineage markers assigned as “other”. The relative abundance of all major lineages was determined out of total cells per FOV and the relative frequency of cell subsets were determined out of total cells of a given lineage, per FOV.

### Region Masking

Region masks were generated to define histologic regions of each FOV including the epithelium, stroma, myoepithelial (periductal) zone, and duct, which was further subdivided into the duct edge, duct mid, and duct core. We removed gold-positive area which marked regions of bare slide from holes in the tissue, providing an accurate measurement of tissue area. This area measurement could be used to calculate cellular density in specific histologic regions, e.g., fibroblast density in the stroma, which was critical to normalize the observed cell abundances by how much tissue of a specific type was sampled, and prevent bias based on how much tumor vs stroma the FOV covered. The epithelial mask was first generated though merging ECAD and PanCK signal and applying smoothing and radial expansion to incorporate the myoepithelial zone, and the inside of ducts were filled. The stromal mask included all image area outside of the epithelial mask. Duct masks were generated through the erosion of the epithelial masks by 25 pixels. The myoepithelial mask was generated by subtracting the duct mask from the epithelial mask. Duct edge, duct mid, and duct core masks (Figure 3I) were generated by eroding the duct mask by subsequent 100-pixel increments.

### Protein and Cellular Spatial Enrichment Analyses

A spatial enrichment approached was used as previously described (Keren et al., 2018, 2019; McCaffrey et al., 2020) to identify patterns of protein enrichment or exclusion across all protein pairs. HH3 was excluded from the analysis. For each pair of markers, X and Y, the number of times cells positive (normalized expression >0.25) for protein X was within a ~50 um radius of cells positive for protein Y was counted. A null distribution was produced by performing 100 bootstrap permutations where the locations of cells positive for protein Y were randomized. A z-score was calculated comparing the number of true cooccurrences of cells positive for protein X and Y relative to the null distribution. Importantly, symmetry is assumed: the values of when calculating the spatial enrichment of protein X close to protein Y are the same as with protein Y close to protein X. For each pair of proteins X and Y the average z-score was calculated across all DCIS FOVs.

To analyze cellular associations with the myoepithelium, the distances between all cell centroids to the nearest perimeter location of the myoepithelium mask (described above) were calculated. To quantity cell type spatial interactions, the mean distances between cell centroids for all cell phenotype pairs (self-self pairs excluded) were calculated per region.

Cell neighborhoods were produced by first generating a cell neighbor matrix, where each row represents an index cell, and the columns indicate the relative frequency of each cell phenotype within an 36um radius of the index cell. Next the neighbor matrix was clustered to 10 clusters using k-means clustering. Neighborhood cellular profile was determined by assessing the mean prevalence of each cell phenotype in the index cells’ 36um radius, while functional marker expression was determined by assessing mean marker expression by the index cells assigned to each neighborhood cluster.

### DCIS UMAP Visualization

UMAP embeddings were determined for all DCIS tumors (pure, synchronous, primary and recurrent) using the R implementation (McInnes et al., 2020) with the following parameters: n_neighbors = 15, min dist = 0.1 and the following markers: PanCK, CK7, CK5, ECAD, VIM, ER, HER2, AR, CD31, SMA, CD45, HLADR, CD68, CD11c, CD14, CD20, CD3, CD4, CD8, MPO, Tryptase.

### EMT GSEA

To identify genes and pathways associated to EMT, MIBI-identified DCIS vimentin high vs low samples were selected, and the epithelial fraction of an adjacent tissue section was analyzed by LCM-RNAseq (Vim high, n = 26; Vim low, n = 32). DESeq2 R package (version 1.30.0) was used for data normalization and differential expression analysis. Results were sorted by decreasing log fold change and the ranked list was subjected to GSEA against C2 curated dataset of molecular signature database (MSigDB)(Subramanian et al., 2005). P values were corrected for multiple comparisons by using Benjamini-Hochberg method and terms with p adj < 0.05 were considered.

### ECM Gene Analysis

To analyze extracellular matrix components by gene expression, an extracellular matrix gene signature (GO extracellular matrix structural constituent, GO:0030021) was downloaded from GSEA website and used to compare MIBI-identified samples with the top and bottom quartiles of cancer associated fibroblast density in the stroma. Stromal LCM-RNAseq samples were used for this analysis. Raw reads were normalized with DESeq2 R package (version 1.30.0)(Anders and Huber, 2010) and a paired T-test was compared to the log2 ratio of group means to generate the volcano plot.

### Myoepithelial Continuity and Thickness Analysis

To define a window of myoepithelial signal quantitation, we used a topology-preserving operation to define a curve 5 pixels out from the epithelial mask edge (see *Region Masking*) and a curve 30 pixels in from the epithelium mask edge, and we defined those pixels in between these two curves as the myoepithelium mask. We subdivided the outer curve into 5-pixel long arc-segments, and for each point on the outer edge in between two segments, found the nearest point on the inner edge, dividing the myoepithelium into a string of quadrilaterals or “wedges”. Wedges are then subdivided each wedge along the in-out (of the epithelium) axis into 10 segments. Wedges are merged when both their combined inner and outer edges has an arc-length less than 15 pixels.

We took pre-processed (background subtracted, de-noised) SMA pixels within the mesh and smoothed them with a Gaussian blur of radius of 1. We then calculated the density of SMA signal within each mesh-segment as the mean pixel value of smoothed SMA within that mesh-segment. This density was then binarized to create a SMA-positivity mesh, using a threshold of 0.5 (density > 0.5 as positive).

The percentage of duct perimeter covered by myoepithelium was calculated by assigning an “SMA-present” variable to each wedge, “0” if no mesh-segments in the wedge were positive for SMA, and “1” otherwise. Each wedge is weighted by its area relative to the myoepithelium area. The sum over all wedges of the product of the “SMA-present” variable and the weight was defined as the percent perimeter SMA positivity. The average (non-zero) thickness of the myoepithelium for each duct was calculated by finding the weighted average “wedge thickness” for SMA-positive wedges (”SMA-present” was 1). The wedge thickness was calculated as the distance between the inner-most and outer-most positive mesh-segments. The positive wedges were weighted by their area relative to the total area of positive wedges.

The percent myoepithelial-covered perimeter and average myoepithelial thickness metrics were waited over meshes (ducts) in a given image by assigning a weight to each duct equal to the total area of the duct myoepithelium divided by the sum of the total areas of all myoepithelium in the image that met a minimum size filter of 7500 pixels.

### Myoepithelial Pixel Clustering Analysis

Pre-processed (background subtracted, de-noised) images were first subset for pixels within the myoepithelium mask. Pixels within the myoepithelium mask were then further subset for pixels with SMA expression greater than 0. For all SMA^+^ pixels within the myoepithelium mask, a Gaussian blur was applied using a standard deviation of 1.5 for the Gaussian kernel. Pixels were normalized by their total expression, such that the total expression of each pixel was equal to 1. A 99.9% normalization was applied for each marker. Pixels were clustered into 100 clusters using FlowSOM (Van Gassen et al., 2015) based on the expression of 6 markers: PanCK, CK5, Vimentin, ECAD, CD44, and CK7. The average expression of each of the 100 pixel clusters was found and the z-score for each marker across the 100 pixel clusters was computed. All z-scores were capped at 3, such that the maximum z-score was 3. Using these z-scored expression values, the 100 pixel clusters were hierarchically clustered using Euclidean distance into 6 metaclusters. SMA^+^ pixels that were negative for the 6 markers used for FlowSOM were annotated as the SMA-only metacluster, resulting in a total of 7 metaclusters. These metaclusters were mapped back to the original images to generate overlay images colored by pixel metacluster.

### Collagen Morphometrics

To identify collagen fibers the background-removed Col1 images are first preprocessed: Col1 pixel intensities were capped at 5 and gamma transformed (1 of 2), and contrast enhanced. Images are then blurred via gaussian with sigma of 2. While this enhances fidelity, it gives less clear ‘0-borders’. This is mitigated by generating a ‘0-region’ mask and setting all values to 0 in that region. Then, highly localized contrast enhancement is applied. Raw fiber signal intensity can vary greatly within a FOV, so this step helps to enhance locally recognizable, but globally dim fiber candidates. After this process, contrast is globally enhanced via a reverse gamma transformation (2 of 2).

Collagen fiber objects are generated by watershed segmentation on the preprocessed images. An adaptive thresholding method was developed to appreciate variability in total image intensities across the large dataset. A dilated and eroded version of each preprocessed image was produced and subjected to multiotsu thresholding. For thin fibers, the higher watershed region is set to everywhere where the eroded image has greater intensity than the highest multiotsu threshold for the eroded image, while the lower watershed region is set to everywhere where the dilated image has lower intensity than the highest multiotsu threshold for the eroded image. For thick fibers, the same procedure is performed, except the lower watershed region uses the middle multiotsu threshold for the dilated image. Elevation maps for watershed are generated via the sobel gradient of a blurred version of the preprocessed images. Once objects are extracted and segmented, length, global orientation, perimeter, and width are computed for each object. Objects which cover low intensity regions of the image are treated as preprocessing artifacts and are not included in averaging.

For fiber alignment scoring, fibers are filtered for elongated shape (length > 2*width), and alignment is scored as the normalized total paired square difference over its k nearest neighbors (k = 4 was chosen). To accommodate for the elongated shape of these object, K-nearest neighbors were computed with the ‘ellipsoidal membrane distance’ (EM distance), which is the Euclidean centroid distance minus the portion of said distance that lies within the ellipse representation of the object.

### Cibersort Analysis

CIBERSORTx (CSx)(Newman et al., 2019) was used to infer the immune fraction in LCM-SMART3SEQ samples. We first generated a tissue resident immune cell signature matrix by using a published breast cancer scRNAseq dataset, downloaded from Gene Expression Omnibus database (GEO data repository accession numbers GSE114727, GSE114725)(Barrett et al., 2013). Normalized counts were obtained by using Seurat R package (version 3.2.0). The resultant signature matrix contained 3484 genes and allowed to resolve different immune cell types, including B, CD8 T, CD4 T, NKT, NK, mast cells, neutrophils, monocytes, macrophages and dendritic cells. The signature matrix was first in-silico validated. In order to test the accuracy of the signature matrix, a set of samples from the same scRNAseq dataset was reserved to build a synthetic matrix of bulk RNAseq data. By mixing different proportion of single cells transcripts, the synthetic bulk was used to analyze the correlation between known vs obtained cell proportions by CSx. Pearson’s coefficient was above 0.75 in all of the cases, most of them above 0.9. Therefore, we used the aforementioned matrix to deconvolve the LCM-RNAseq samples and to compare CSx-estimated cell abundance with MIBI-identified cell types.

### Prediction of recurrence

To predict recurrence, we identified patients in the cohort with follow-up data demonstrating carcinoma recurrence (n=12), invasive recurrence (n=19), or at least 11 years without recurrence (n=47). For each patient, a vector of summary statistics was generated from MIBI data using only images derived from the original lesion. The cohort was split into training and test sets (80/20%); all model optimization and predictor selection used only the training set. Any missing values were replaced with the set’s predictor mean. Predictors with <12 unique values in the training set were dropped from the analysis. Two-class random forest probability models (ranger package)(Wright and Ziegler, 2017) were trained to discriminate recurrence versus non-recurrence, and invasive recurrence versus non-recurrence. Hyperparameters were tuned to minimize out-of-bag error. One tuned hyperparameter was predictor subset selection by correlation thresholding: predictors were ranked in importance by performing a KS test between recurrence and non-recurrence. Greater importance was placed on predictors with lower p-values, with ties broken by weighting predictors with greater coefficients of variance (CV). All predictors were correlated (Spearman method) and correlations were thresholded (*invasive* r>|0.5|, *all recurrence* r>|0.6|). For each group of correlated predictors above a given threshold, only the highest-ranked predictor was used in the model. The optimized random forest model was evaluated on the test set and a receiver operating characteristic (ROC) curve was generated (pROC package)(Robin et al., 2011) using the model’s assigned probability scores. Area under the curve (AUC) was calculated with 95% confidence intervals, determined by bootstrapping. Each predictor’s importance was evaluated in the model by its Gini index. Similarly, two-class random forest probability models were also trained using only clinical parameters as predictors (age, mammograph density, tumor grade, and tumor necrosis) without subset selection. For the MIBI-based predictions, an optimal probability threshold was selected by the Youden method to assign predicted class to the test set, and Kaplan-Meier curves were calculated (survival package)(Therneau and Grambsch, 2000).

### Statistical Analysis

All statistical analyses were performed using GraphPad Prism software or in R. Grouped data is presented with individual sample points throughout, and where not applicable, data is presented as a mean with standard deviation. For determining significance, grouped data was first tested for normality with the D’Agostino & Pearson omnibus normality test. Normally distributed data was compared between two groups with the two-tailed Student’s T-test. Non-normal data was compared between two groups using the Mann–Whitney Test. Multiple groups were compared using the Dunn’s Multiple Comparison Test.

### Software

Image processing was conducted with Matlab 2016a and Matlab 2019b. Statistical analysis was conducted in Graphpad Prism. Data visualization and plots were generated in R with ggplot and pheatmap packages, in Graphpad Prism, and in Python using the scikitimage, matplotlib, and seaborn packages. Representative images were processed in Adobe Photoshop. Schematic visualizations were produced with Biorender. R packages for GSEA: AnnotationDbi, 1.52.0 & org.Hs.eg.db, 3.12.0, clusterProfiler, version 3.19.0, for GSEA msigdbr, version ‘7.2.1’, for C2 curated datasets. Python packages for spatial enrichment analysis and collagen morphometrics: sckikit-image, pandas, numpy, xarray, scipy, statsmodels.

## Data and Code Availability

All custom code used to analyze data will be made available through a Github repository and all processed images and annotated single cell data will be made available on a Human Tumor Atlas Network public repository.

## Author Contributions

TR conceived the study design, performed experiments, analyzed data, and wrote the manuscript with MA. DG developed the classifier model and performed related analyses. CCL developed the myoepithelial pixel clustering approach and performed related analyses. SHS. processed the LCM-RNAseq data and BRG performed all RNAseq analyses. EFM assisted with data analysis with AK, LK, and SV. N.F.G. assisted with image segmentation. AB developed and performed the myoepithelial morphology analyses and AK performed the collagen morphology analyses. GAC, DJV, KD assisted with cohort design and patient sample preparation, and SS performed pathological review, and SV and ZK assisted with immunohistochemistry. SEH, SCB, RBW and MA supervised the work.

## Acknowledgements

The authors thank the HTAN Consortium for the intellectual and collaborative support of this work. We would like to thank Pauline Chu and the Stanford Human Histology Core for providing technical assistance. TR was supported by the American Cancer Society Postdoctoral Fellowship 133099-PF-19-002-01-CCE, and Stanford Immunology Training Grant 5 T32 AI07290-33. DRG was supported by the Bio-X Stanford Interdisciplinary Graduate Fellowship. CCL was supported by the Stanford Graduate Fellowship. RBW was supported by R01CA193694 and U2C CA233254. MA was supported by 1-DP5-OD019822. SCB and MA were jointly supported by 1R01AG056287 and 1R01AG057915, 1U24CA224309, the Bill and Melinda Gates Foundation, and a Translational Research Award from the Stanford Cancer Institute.

