## Supplemental Figures and Legends for "Transition to invasive breast cancer is associated with progressive changes in the structure and composition of tumor stroma"

### Supplemental Figure Legends

#### Table S1. DCIS cohort patient information

Clinical, demographic, and technical information about each patient tumor included in the study cohort.

#### Table S2. MIBI-TOF antibody panel

Antibodies used in imaging panel. Direct metal conjugates are listed with the clone, metal isotope, and titer used in the study, with the overnight and 1hr staining panels separately listed. Information of primary hapten-conjugated antibodies are also included. Low-level image processing information for each marker is listed to the right including signal quantitation window (*start, stop*) and denoising and aggregate-removal thresholds.

#### Table S3. Related to Figure 7

Tumor features selected by the different classifier models are listed with accompanying stats, ranked by Gini importance.

#### Figure S1. Related to Figure 1

Representative immune marker staining in control tissues tonsil, lymph node, and placenta.

#### Figure S2. Related to Figure 2

**A**. Workflow for Deepcell-based segmentation of single cells from multiplexed images. **B**. Conceptual overview of hierarchical FlowSOM algorithm application.

#### Figure S3. Related to Figure 3

**A**. Criteria used to define tumors as ER, AR, HER2, or Ki67 positive, and HER2-intense. **B**. Histogram of single cell HER2 normalized expression levels in normal breast (Norm.), DCIS, synchronous tumors (Sync.) and IBC. Cutoffs for HER2 medium and intense expression are shown. **C**. Boxplots showing the distribution of HER2-medium or HER2-intense tumors in each progression group. **D**. GSEA results from the C2 database of genesets significantly enriched in DCIS tumors that are VIM-high (left side) or VIM-low (right side). Normalized enrichment score (NES) is denoted by the point size, adjusted p value is denoted by color.

#### Figure S4. Related to Figure 5

**A**. Dotplot of CAF frequency in tumors with Ki67$^+$ fibroblasts. **B.** Dotplot of CAF frequency in tumors with >10 pS6$^+$ cells.

#### Figure S5. Related to Figure 6

**A**. Violin plot showing immune lineage frequency of total cells, in the stroma and duct, as detected by MIBI imaging or Cibersortx estimated fraction. **B**. The mean density of each immune cell type is shown in the stroma and duct in normal breast (N), DCIS (D), synchronous tumors (S), and IBC (I). **C.** Heatmaps showing the z-score normalized mean expression for functional markers in each cellular neighborhood in macrophages. **D.** Cellular neighborhood map with a zoomed ROI of a LyE1-enriched region showing a MIBI overlay PDL1 expression and macrophage and T cell markers.

**Figure S6. Related to Figure 7**

A. Stacked bar plot showing the frequency of mastectomy, radiation therapy, and tamoxifen therapy in the case and ctrl outcome groups in the training data for the invasive recurrence model (top) and all-recurrence model (bottom). **B.** The distribution of mastectomy, radiation, and tamoxifen therapy is shown by color in the model-predicted cases and ctrls, with the random forest prediction probability shown for each patient. P values comparing the treated frequency of total between groups is displayed. Invasive recurrence model (left) and all recurrence model (right) shown separately. **C.** MIBI features with top classifier importance for the all-recurrence model are shown, ranked by Gini importance. Features are colored based on enrichment either in cases (orange) or controls (green), importance bars are colored based on the feature utilizing spatial information (purple) or not (gold). **D.** The distribution of spatial vs non-spatial features are shown for all features identified (total), those used by the model (selected), and those in the top 20 most important features (top 20). **E**. Predicted survival of patients identified in the test set of the invasive-recurrence model as case or control. **F.** Stacked bar plot showing the frequency of features from each cellular lineage or ECM, compared between the total training set, those selected by the classifier models, and the top 20 important features from each model.

Figure S1. Related to Figure 1

Tonsil

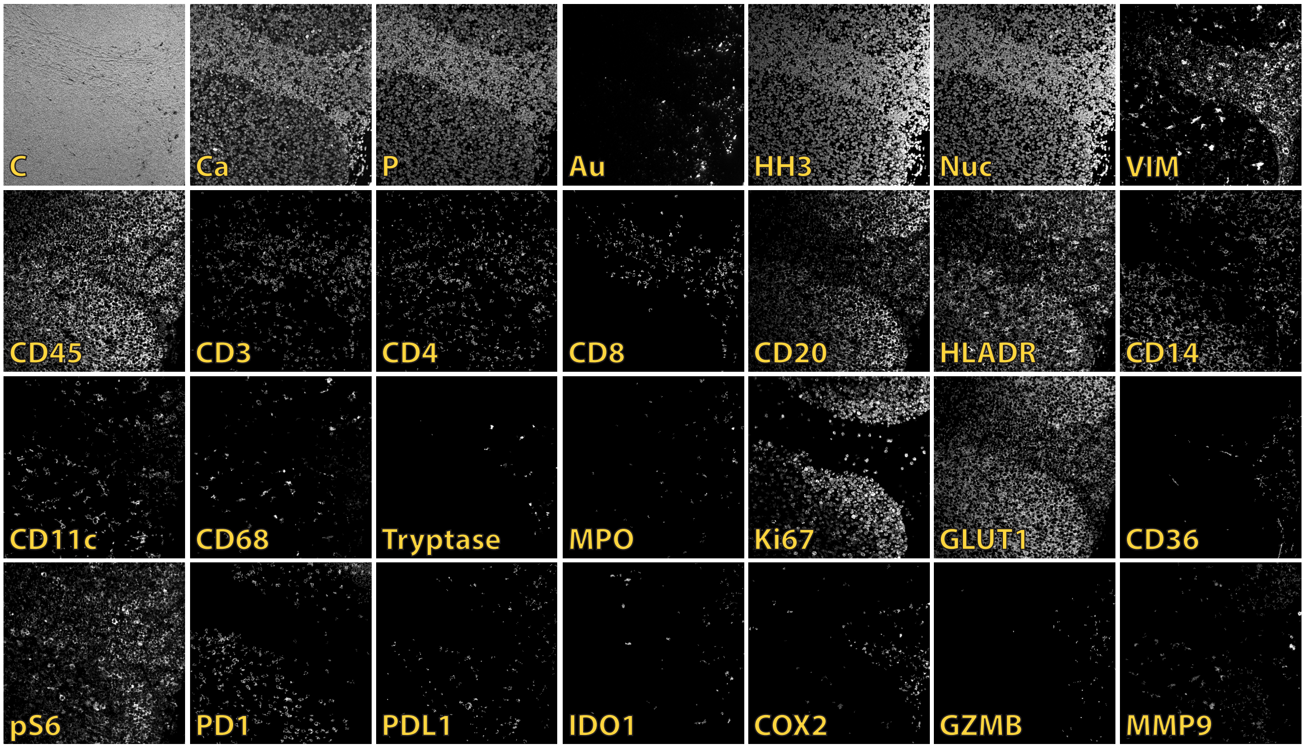


Lymph Node

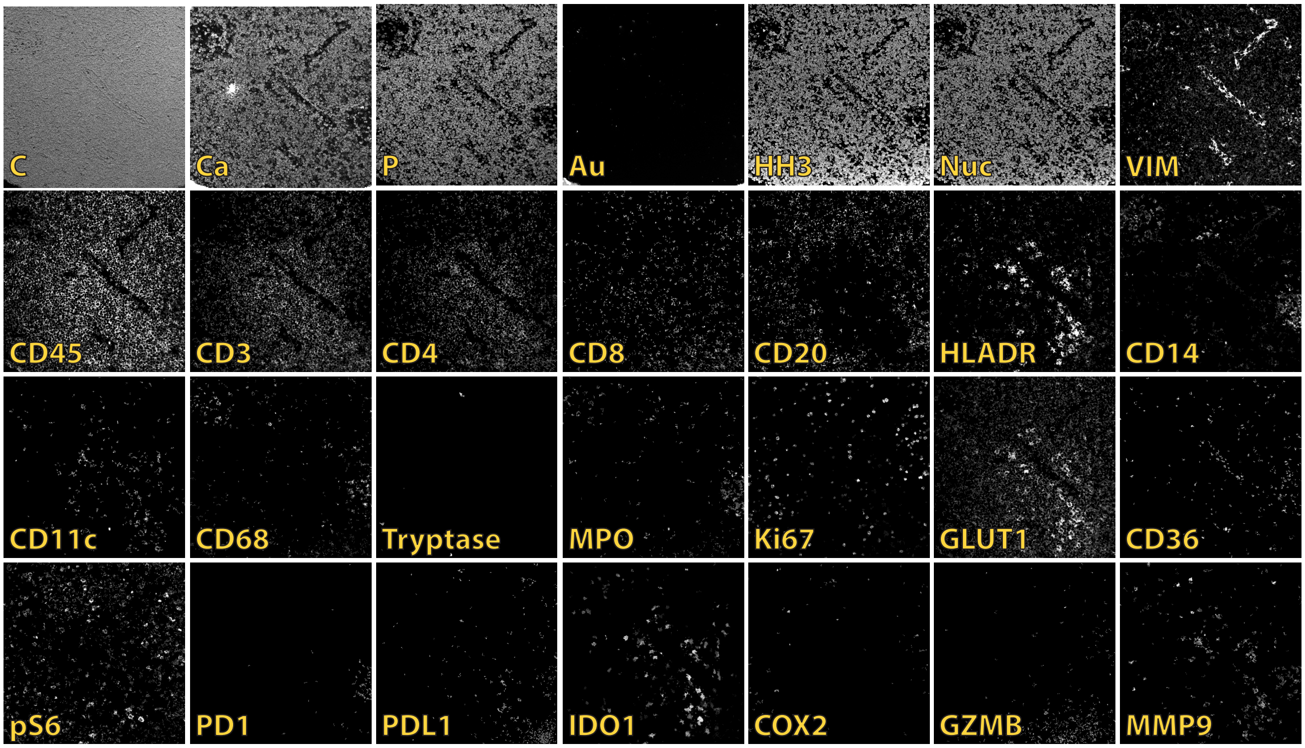


Placenta

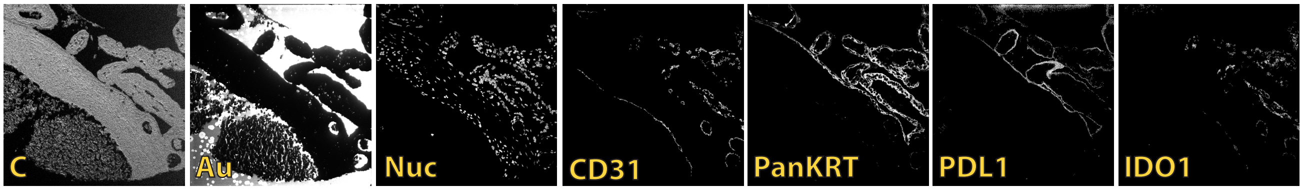

Figure S2. Related to Figure 2

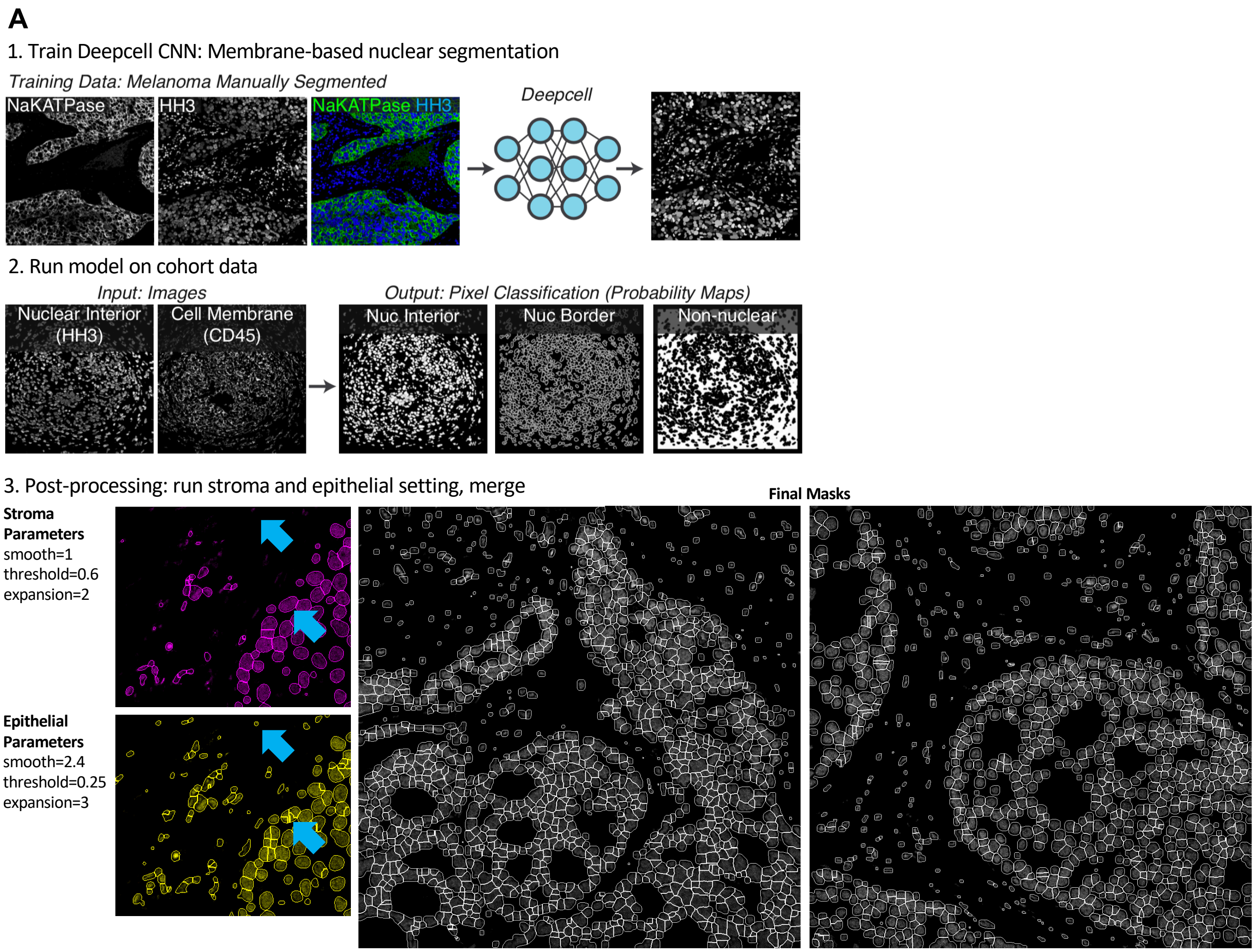


**B** FlowSOM lineage clustering strategy

**Round 1: Lineages** / **Round 2: Lineage Subsets**

- All cells
  - Tumor
    - Luminal (CK7+, ECAD+, PanCK+)
    - Basal (CK5+, ECAD+, PanCK+)
    - EMT (VIM+, ECAD+/-, PanCK+/-)
    - CK5/7low (ECAD+, PanCK+)
  - Myoepithelial
  - Immune
    - Monocyte (CD14+, HLADR+/-, CD45+)
    - MonoDC (CD14+, CD11c+, HLADR+, CD45+)
    - DC (CD11c+, HLADR+, CD45+)
    - Macs (CD68+, HLADR+, CD14+/-, CD4+/-, CD45+)
    - APC (HLADR+, CD45+)
    - Mast (Tryptase+, CD45+)
    - Neut (MPO+, CD45+)
    - CD4 T (CD4+, CD3+, CD45+)
    - CD8 T (CD8+, CD3+, CD45+)
    - B cell (CD20+, HLADR+/-, CD45+)
    - ImmOther (CD45+)
  - Endothelial
  - Fibroblast
    - Normal (CD36+, VIM+)
    - Myo (SMA+, VIM+)
    - CAF (FAP+, SMA+/-, VIM+)
    - Resting (VIM+)
  - Other

Figure S3. Related to Figure 3

**A**

| Marker | Marker Threshold | Tumor Positivity Criteria |
|---|---|---|
| ER | <0.25 | >1% of Tumor Cells Positive |
| AR | <0.25 | >1% of Tumor Cells Positive |
| HER2 | <0.2 pos, 0.7 intense | >10% of Tumor Cells Intense |
| Ki67 | <0.25 | >14% of Tumor Cells Positive |

**B**

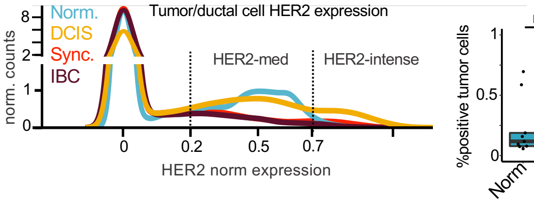


**C**

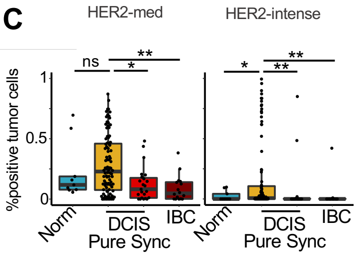


**D**

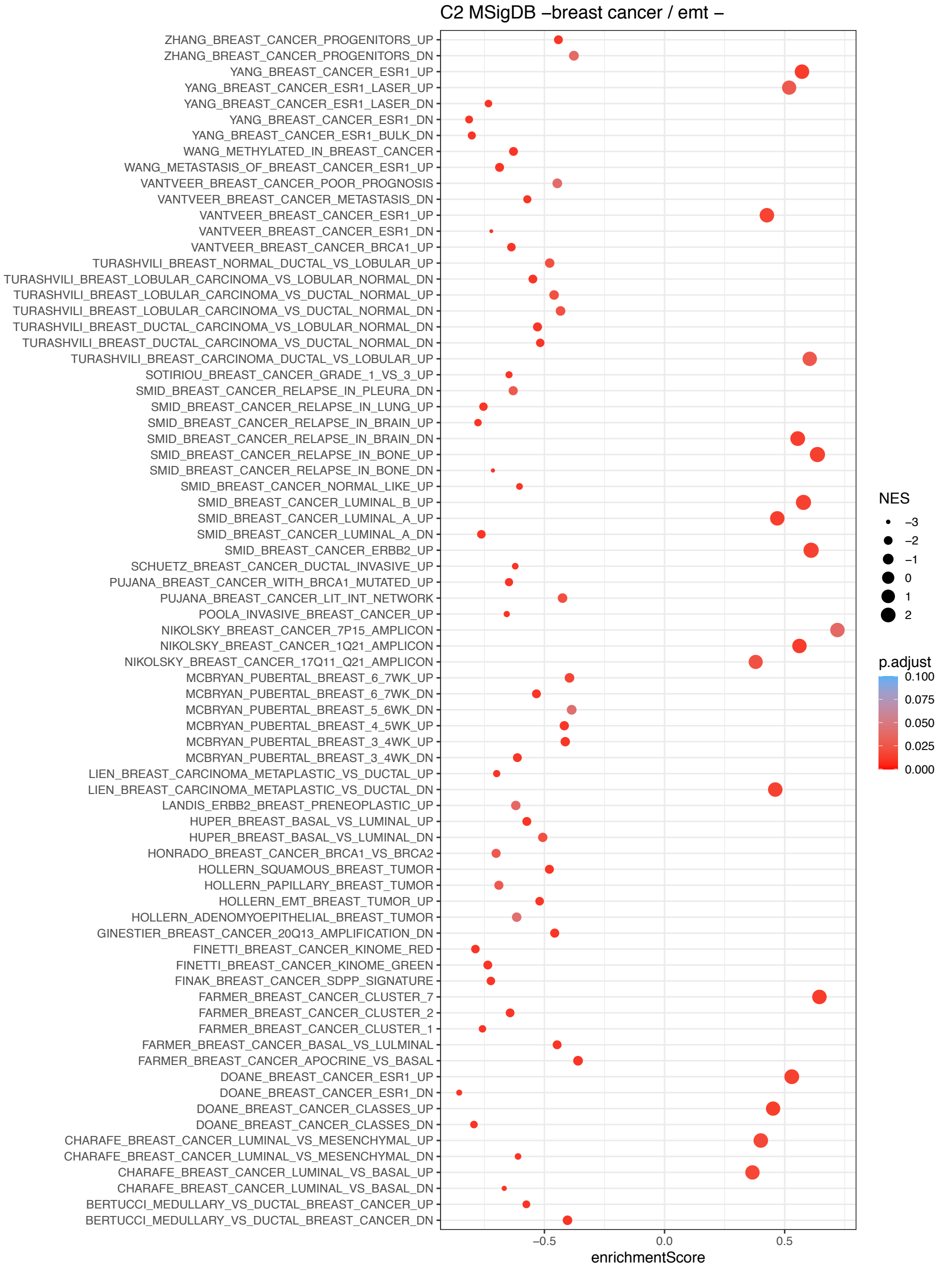

Figure S4. Related to Figure 5

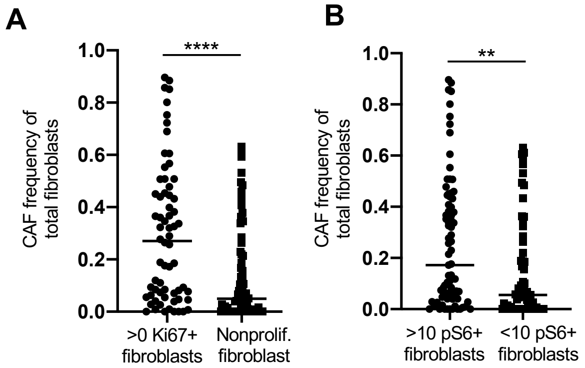

Figure S5. Related to Figure 6

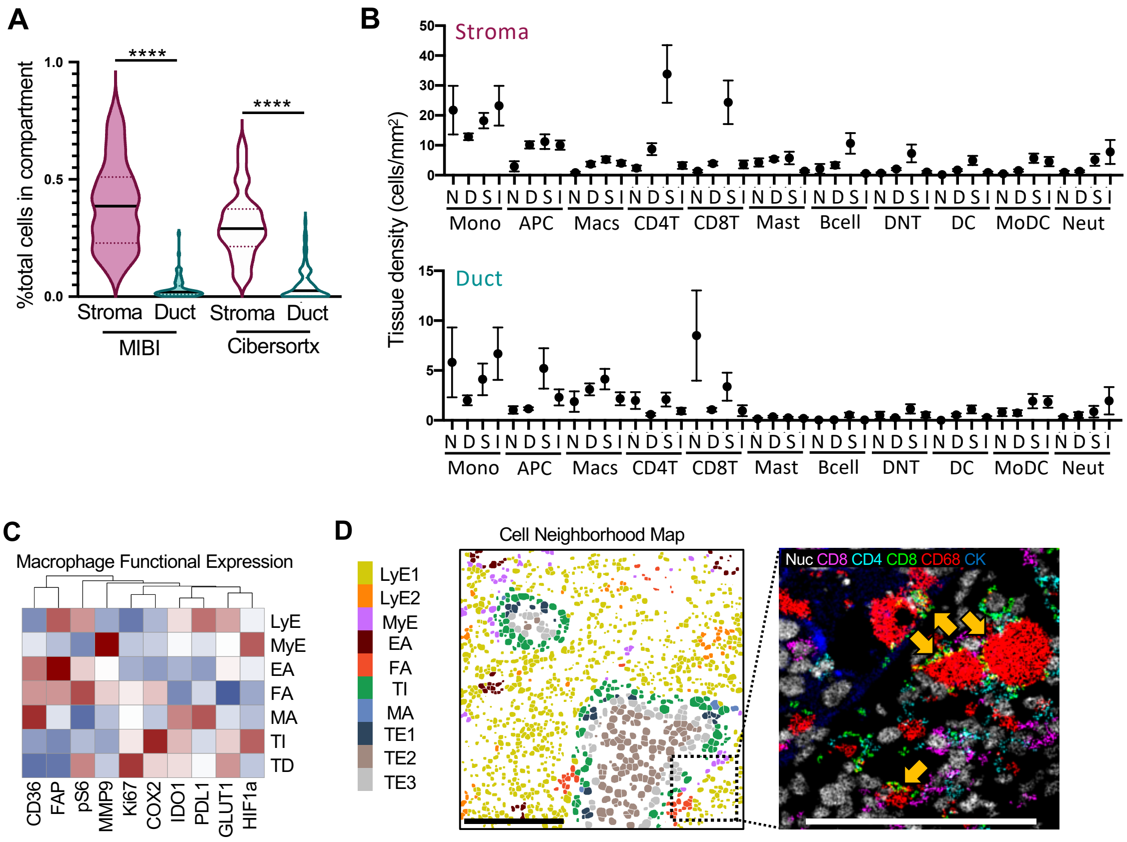

Figure S6. Related to Figure 7

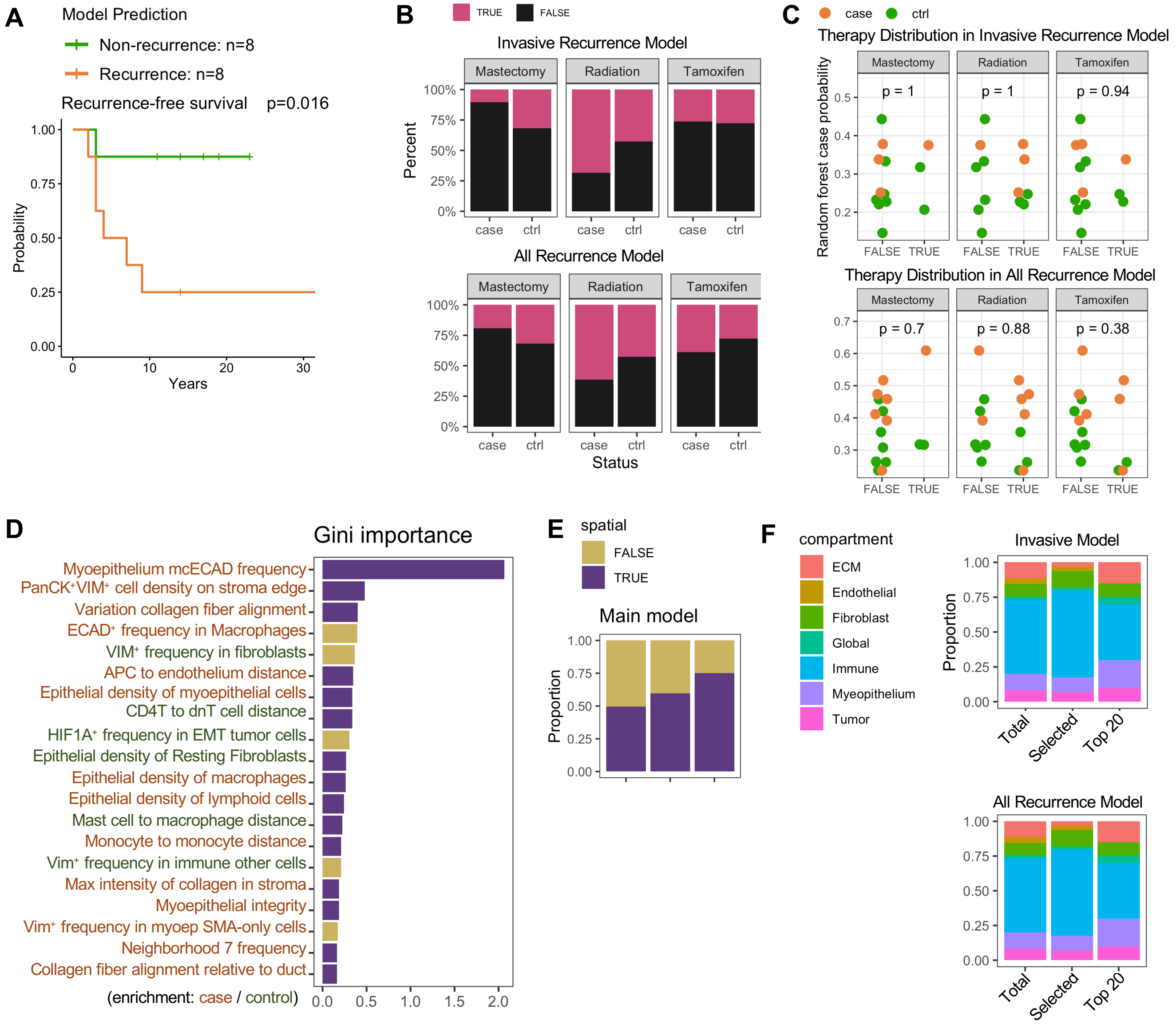
